# Learning cell communication from spatial graphs of cells

**DOI:** 10.1101/2021.07.11.451750

**Authors:** David S. Fischer, Anna C. Schaar, Fabian J. Theis

## Abstract

Tissue niches are sources of cellular variation and key to understanding both single-cell and tissue phenotypes. The interaction of a cell with its niche can be described through cell communication events. These events cannot be directly observed in molecular profiling assays of single cells and have to be inferred. However, computational models of cell communication and variance attribution defined on data from dissociated tissues suffer from multiple limitations with respect to their ability to define and to identify communication events. We address these limitations using spatial molecular profiling data with node-centric expression modeling (NCEM), a computational method based on graph neural networks which reconciles variance attribution and communication modeling in a single model of tissue niches. We use these models in varying complexity across spatial assays, such as immunohistochemistry and MERFISH, and biological systems to demonstrate that the statistical cell–cell dependencies discovered by NCEM are plausible signatures of known molecular processes underlying cell communication. We identify principles of tissue organisation as cell communication events across multiple datasets using interpretation mechanisms. In the primary motor cortex, we found gene expression variation that is due to niche composition variation across cortical depth. Using the same approach, we also identified niche-dependent cell state variation in CD8 T cells from inflamed colon and colorectal cancer. Finally, we show that NCEMs can be extended to mixed models of explicit cell communication events and latent intrinsic sources of variation in conditional variational autoencoders to yield holistic models of cellular variation in spatial molecular profiling data. Altogether, this graphical model of cellular niches is a step towards understanding emergent tissue phenotypes.

---

Cells interact on multiple length-scales through direct contact of surface-bound receptors and ligands, tight-junctions and mechanical effects, as well through indirect mechanisms, including soluble factors and vesicles^1^. These communication events can usually not be directly observed but are critical to understand emergent phenomena in tissue niches. However, the sender and receiver cell of a communication event are often characterized by molecular signatures, both facilitating the signalling, such as ligand and receptor expression^2–4^, and resulting from the signalling event, such as intracellular cascades induced by receptor activation. These molecular signatures are used in computational methods to infer latent cell communication events in a tissue. Core assumptions inherent in these algorithms can be summarized in two groups: First, the co-occurrence of ligand and receptor expression across cell-types is used in multiple computational models to suggest putative axes of cell communication^2,3^. Second, from a statistical point of view, cell communication is a source of cellular variance. The gene expression of a receiving cell depends on the sending cells in the local tissue niche, thus inducing a statistical dependency that can be used to infer communication events^2,5^. Here, we address three core issues inherent in cell communication inference methods founded on these two assumptions using spatial information. First, axes of cell communication cannot be necessarily disentangled based on data from dissociated cells because of the large number of potential interactions in a population of cells. We propose to use spatial information from image-structured molecular profiling assays to constrain the inference of such gene expression dependencies between neighboring cells. For this purpose, we define a prior distribution on the possibility of a communication event between two cells as a function of their distance, using space to prioritize pairs of cells for which communication is molecularly plausible. Second, cell pairs with matching ligand and receptor expression can occur in a tissue even in the absence of communication, for example because of spatial separation in the tissue architecture. We propose to use the receiver cell molecular signature as conclusive evidence for the presence of a communication event. Third, molecular signatures of receiver cells can be observed independent of ligand–receptor quantification. Therefore, cell communication inference should be possible both in the absence of a reliable quantification of ligand or receptor genes, which occurs in targeted molecular profiling assays, or in cases in which these genes are expressed at low levels, and in receptor protein-free communication, such as in physical interactions. We define a generalized framework to detect receiver cell signatures detached from ligand–receptor definitions.

We reconcile these assumptions in a statistical model for cell communication events in spatial molecular profiling data, referred to as a node-centric expression model (NCEM). NCEMs are trained on segmented image-structured molecular data from assays with subcellular resolution, such as immunohistochemistry^6,7^, imaging mass cytometry^8^, and MERFISH^9^ (Fig. 1a). We enforced a neighborhood constraint on communication events using spatial graphs of cells, where the graph serves as prior for cellular communication. The resulting model is a graph neural network^10,11^ and constitutes an extension of models of dissociation-based data, in which cells are modeled independently, to a spatially-constrained model of cell–cell dependencies (Fig. 1a). This model relies on the stratification of cells into clusters that can then be treated as categorical sender and receiver cell-types in communication events. Importantly, the directionality of these sender–receiver signalling is defined on the level of pairs of single cells by the statistical association of the molecular state of the receiver on the sender. We discovered these statistical associations in a model of the molecular state of cells conditioned on the putative sender cell-types present in their respective spatial neighborhoods. The complexity of the spatial dependencies observed in a tissue strongly depends on the tissue architecture and the ability of the molecular readout to capture the signatures of these dependendencies. Therefore, the cell communication model complexity needs to be adapted to the scenario at hand. We studied three increasingly complex models of cell–cell dependencies. First, we propose a linear graph neural network that can be framed as a graph-aware generalized linear model (GLM) to model linear expression effects of cell communication. Secondly, we generalize NCEMs to nonlinear graph neural networks that can account for higher-order cell interactions. Third, we consider generative latent variable models that also model confounding latent sources of variation, such as cell intrinsic effects. We interpret NCEM fits in terms of cell communication patterns. We demonstrate cell communication inference with NCEMs on five datasets (Online methods, Supp. Figure 2): a MERFISH dataset of mouse brain^12^ (MERFISH – brain) of 634 images across with 254 genes observed in 284,098 cells, a chip cytometry dataset of an inflamed colon (Data Availability) (chip cytometry – colon) of two images with 19 genes observed in 11,321 cells, a MIBI TOF dataset of colorectal carcinoma^8^ (MIBI TOF – cancer) of 58 images with 36 genes observed in 63,747 cells, a MELC dataset of tonsils^7^ (MELC – tonsils) of one image with 51 genes observed in 6,991 cells and a CODEX dataset of colorectal cancer^13^ (CODEX – cancer) of 140 images with 57 genes observed in 272,266 cells. We discover cell–cell dependencies at molecularly plausible length scales and attribute molecular variation within cell-types to niche composition. The inferred interactions between cells serve as a powerful mechanism to interpret niche composition variation and its effect on the contained cell.

**Figure 1:**
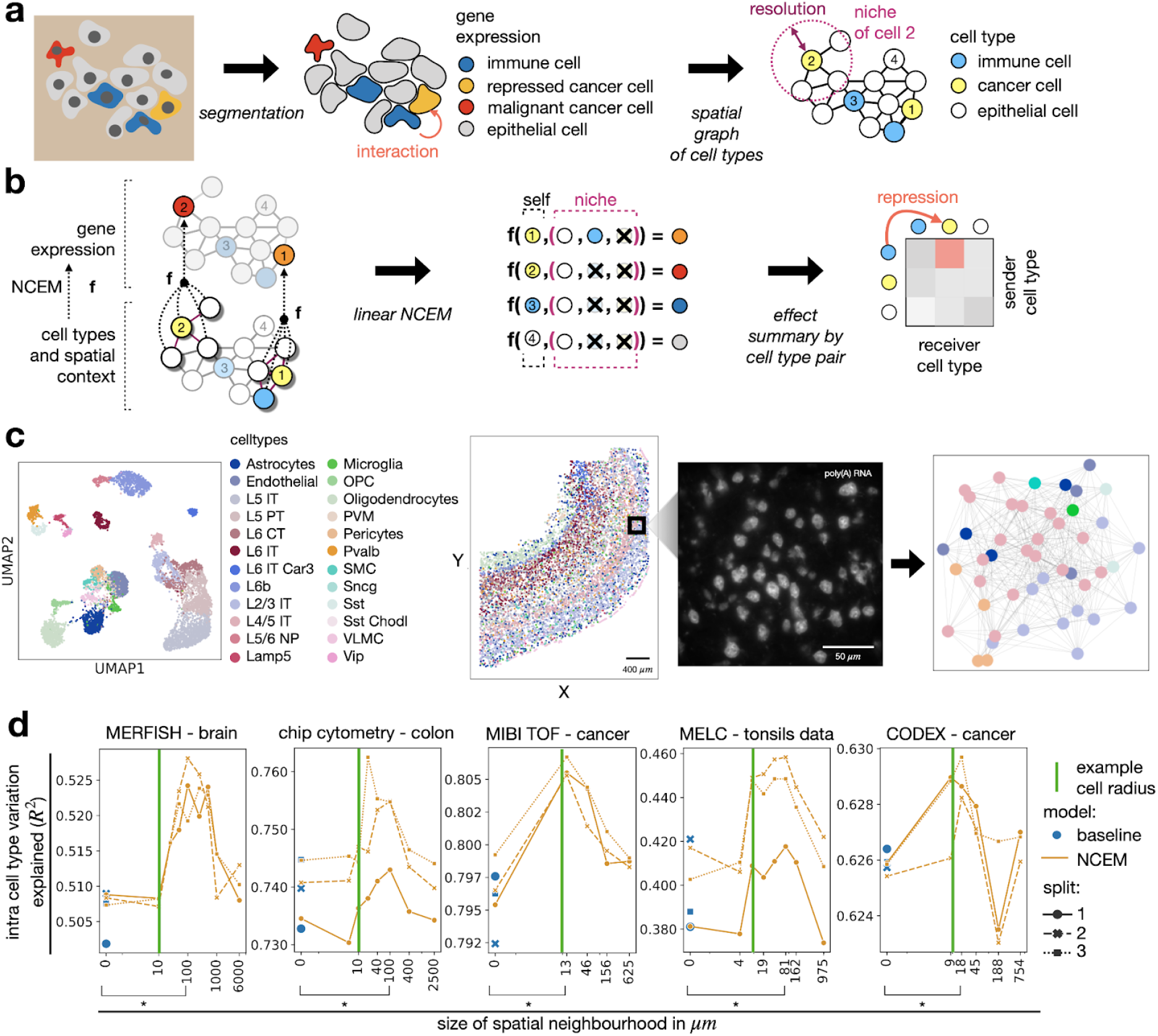
Modeling cell communication as spatial cell state dependencies. **(a)** Spatial graphs of cells can be computed from spatial molecular profiling data. After segmentation of cells or nuclei, each cell is characterized by a vector of gene expression measurements which are coarse-grained in a clustering to categorical cell-types. A graph can be assembled based on proximity of segments. *resolution*: radius of circle used to define niche. **(b)** Node-centric expression models (NCEMs) describe the gene expression of a cell (color of the node) as a function of its spatial neighborhood (niche). This function is a graph neural network and can be reduced to a linear model in the simplest case. This linear model contains directional effects from sender to receiver cell-types which can be summarised in an effect matrix between cell-types. *Y*: gene expression vector of a cell. **(c)** Proximity graphs in spatial transcriptomics data. Shown are a UMAP of molecular embedding of all cells in slide 153 (n = 7439 cells) with cell-type superimposed, followed by Slice 153 of mouse brain in the MERFISH - brain dataset with the spatial allocation of all cell-types superimposed, field of view number 486 of the same slice with poly(A) RNA channel superimposed at central z-plane (z = 4.5 μm), and the spatial proximity graph of the same field of view with a resolution of 100 μm. **(d)** Linear models capture neighborhood dependencies in spatially-resolved single-cell data. Shown are the R^2^ values between predicted expression vectors and observed expression vectors for held-out test cells of linear models by resolution in μm with cross validation indicated as point shape and line style with relative outperformance of NCEM model versus baseline model in the MERFISH - brain dataset of 1.77%, in the chip cytometry - colon dataset 1.17%, in the MIBI-TOF - cancer dataset 1.04 %, in the MELC - tonsils dataset 4.48% and in the CODEX - cancer dataset 0.28%. *Example cell radius (green line)*: Example length scale of a cell, here chosen as 10 μm; *baseline (blue dot)*: a nonspatial linear model of gene expression per cell-type; *NCEM*: linear model with interaction effects; *bracket (*)*: significant difference in paired t-test between baseline model and best spatial model (MERFISH - brain dataset p = 0.030, chip cytometry - colon dataset p = 0.019, MIBI-TOF – cancer dataset p = 0.018, MELC – tonsils dataset p = 0.029, CODEX – cancer dataset p = 0.026).

## Results

### Node-centric expression models describe cell–cell dependencies on spatial graphs

We infer cell communication from a cell-type-specific gene expression signature that can be predicted from cell-types in the spatial neighborhood. The data type discussed here consists of image-structured data from molecular profiling assays of RNA or proteins, where pixels correspond to tissue slice positions. Each channel contains a molecular abundance readout of a specific gene. To prepare this analysis, we first used segmentation masks to assign pixels in image-structured data to cells (Fig. 1a,c). We then extracted the mean gene expression from these segments to build cell-wise gene expression vectors and clustered these molecular vectors to assign cells to discrete cell-types (Fig. 1a,c). Based on the cell segmentation, we defined the niche of a cell as the set of cells within a circle centred on the cell’s center. The radius of this circle, the “resolution”, is a model hyperparameter (Online Methods, Fig. 1a). We define a node-centric expression model (NCEM) as a function that maps a cell’s type and its spatial context to the cell’s observed gene expression vector (Fig. 1b). This function is a graph neural network in which the node labels are the cell-wise gene expression vectors^10^. Below, we discuss both multilayer graph neural networks and a single-layer model with an indicator graph aggregator function which is equivalent to a linear model (Fig. 1b, Online Methods): The indicator function aggregator summarises a set of one-hot encoded cell-types in a niche to a vector with binary elements that indicate the presence of each cell-type in the niche. This binary embedding is parameter-free and a transformed covariate space with fixed dimensions and thus suitable for a linear model. In addition to this indicator embedding, we also use parametric neighborhood-embeddings for nonlinear NCEMs (Online Methods). The model of a gene expression vector conditioned on a cell-type reflects the assumption that niches modify gene expression states of cell-types, but do not cause cell-type conversions themselves. This conditional model also reflects the assumption that much spatial information is contained in the gene expression profile of a single cell^14^ and that spatial covariates are primarily useful to supervise variance decomposition of gene expression vectors. In summary, the input data to NCEMs consists of three groups: the set of input cell-type labels, the output gene expression features and the spatial proximity graph.

The input cell-type labels group cells to both sender and receiver cell-types, this annotation can be more or less coarse according to the cell-type clustering resolution. Cell-types from one sender class emit a common signal, and cell-types of one receiver class share a common response gene expression signature for a given signal. The coarseness of the sender and receiver cell-type classes serves as an inductive prior which regularizes the model by reducing the number of parameters that model cell communication. For example, an overly fine-grained set of receiver cell-types could attribute niche-dependent cellular variation within a cell-type to between-cluster variation, thus failing to attribute this axis of variation to niche composition.

The identity and number of molecular species measured in a spatial molecular profiling assay vary strongly across protocols and studies. Accordingly, the observable neighborhood-induced gene expression effects vary equally strong. One could include further cell-features related to morphology and molecule distribution in the cell state^15^ to improve the description of the molecular state of a cell to subcellular gene expression variation.

In a spatial proximity graph, cells are connected by edges if their segment centers are not further apart than the resolution of the model^16^ (Fig. 1a). The choice of this resolution depends on the modeled molecular mechanisms of communication, which vary strongly between contact-based and paracrine communication, for example. In this study, we chose multiple resolutions for each dataset in separate analyses. Each resolution is associated with a separate graph that has a specific node degree distribution which depends on the overall tissue architecture. We defined the distances for each dataset such that they cover the range of average node degrees of the given dataset (Supp. Fig. 1b). This screening of resolutions is useful for discovery of statistical dependencies. On the other hand, a fixed, single resolution can be used to test specific hypotheses of cell communication events on a length scale defined by prior knowledge based on molecular mechanisms.

### Linear NCEM identify cell communication on consistent spatial length scales

We model neighborhood-induced cell state changes with a linear graph neural network that predicts the state of a receiver cell conditioned on the presence of putative sender cell-types present in its neighborhood. This model is equivalent in architecture to a graph-aware GLM (Online Methods) and, therefore, represents a convex optimization problem. While limited in their ability to model higher-order cell interactions, such linear models can be readily interpreted in terms of effect sizes, can be used by non-specialists because of their favourable optimization characteristics and allow for correction for confounding factors such as biological conditions.

First, we established the presence of intra-cell-type variance in a variance decomposition on all five datasets (average intra-cell-type variance 40.6%), even in those datasets with very few genes assayed (Supp. Fig. 2, Online Methods). We fit Gaussian GLMs on transformed single-cell expression measurements across multiple neighborhood sizes with four term groups: image covariates, the cell-type, the presence of putative sender cell-types in the neighborhood (the niche), and an interaction term between the cell-type and the niche (Online Methods). As a null model, we included empty neighborhoods and neighborhoods larger than the considered images. We found that these GLMs were most predictive on an intermediate-length scale of around 69 μm in all five datasets (Fig. 1d), showing that cell-cell dependencies indeed only appear on length scales characteristic of direct and short-range molecular mechanisms of cell communication. With further increasing neighborhood sizes, the prediction performance drops again to the level of nonspatial models, indicating that the spatial effect is not simply due to overfitting to images. The NCEMs outperformed non-spatial baseline models by an average Δ R^2^ of 0.0175. This difference was significant in paired t-test between baseline model and best spatial model of (p < 0.05) across five datasets (Fig. 1d). The Δ R^2^ is small compared to the baseline model R^2^ that characterizes between-cell-type variance (0.39 - 0.79) because the cell-type information accounts for a large fraction of the transcriptomic variance in these samples. Between-cell-type variance often dominates variance in single-cell transcriptomics which is a phenomenon that allows for cell-type assignment through clustering algorithms, for example^17–19^. The remaining within-cell-type variance decomposes into technical noise and within-cell-type biological variance. The biological within-cell-type variance can be decomposed into spatial and non-spatial effects. Here, we only model the spatial component, thus explaining the low magnitude of explained within-cell-type variance compared to the total within-cell-type variance. The increased R^2^ achieved by NCEM can be attributed to the niche composition information inherent in the model, thus providing an algorithmic handle on niche biology.

In addition, we also fit GLMs with fewer parameters, without cell-type to niche interaction terms, keeping only non-receiver-specific sender effects, which are less susceptible to overfitting, and also found dependencies on similar length scales (Supp. Fig. 3, Online Methods). We hypothesized that different molecular mediators of cell–cell dependency on different length-scales apply to different cell-types. Indeed, the model performance as a function of length scale by target cell-type varies between cell-types (Supp. Fig. 4).

### Niche composition explains cellular variation

Next, we interpreted the effects learned by the linear NCEMs. To identify neighborhood-induced molecular states in the primary motor cortex, we selected L2/3 IT cells from the MERFISH – brain dataset^12^ based on their strong spatial effects on cell state (Supp. Fig. 4) and their large abundance, with 41,996 cells across the dataset (Supp. Fig. 1a). To identify neighborhood-induced effects on the molecular states without confounding the analysis by between-image batch and condition effects, we clustered and embedded L2/3 IT cells from a single image based on the molecular information alone (Fig. 2a, Online methods). Indeed, the relative prediction performance of the graph-aware GLM at the optimal resolution compared to nonspatial GLM varies across these clusters (Fig. 2b), as measured by the difference of their respective R^2^ on individual cells. Using the presence of different cell-types in the neighborhood as a binary labels vector, we performed a cluster enrichment analysis with Fisher’s exact test on this clustering (Fig. 2c,d, Online Methods) to annotate clusters with enriched neighborhoods: sub cluster 0 is associated with Scng and VLMC, and cluster 4 with L4/5 IT cells, for example (Fig. 2d). We ordered cell-types as putative sender types of L2/3 IT cells by their maximal cluster-wise enrichment p-value. This ordering of cell-types is very different from the ordering obtained from contact frequencies and CellphoneDB^3^ analysis on this image (Kendall rank correlation of −0.04 and −0.12, p-value of 0.79 and 0.45, respectively, Supp. Fig. 6a, 7b, Online methods), highlighting the novel quality of information captured by this approach. Depending on the analysis setting, either L4/5 IT cells and Scng or L4/5 IT cells and VLMC are also discovered as putative partners of L2/3 IT cells by CellphoneDB (Supp. Fig. 7a,b), showing that core associations are reproduced. Next, we established that the L2/3 IT cell sub clusters are spatially localised in distinct areas of the primary motor cortex^12^: Sub cluster 0 is closer to the layer of Scng and VLMC, whereas sub cluster 4 is closer to the layer of L4/5 IT cells (Fig. 2c,d). This spatial patterning becomes apparent when the sub-cluster-wise relative performance spatial and non-spatial model (Fig. 2b) are broken down to cell-wise comparisons and assigned to cells in their spatial context: The spatial model outperforms the non-spatial model strongly on margins of the L2/3 IT layer in the motor cortex, such as in sub cluster 4 of the L2/3 IT cells (Fig. 2e). We repeated this analysis by applying the same models, which were trained on all images, to a second image and found similar associations to neighborhoods, such as significant associations to Scng, VLMC, and L4/5 IT cells (Supp. Fig. 5b-f). In summary, this analysis attributes molecular variation of single cells to their niche and identifies putative sender cell-types which are in line with the overall tissue architecture.

**Figure 2:**
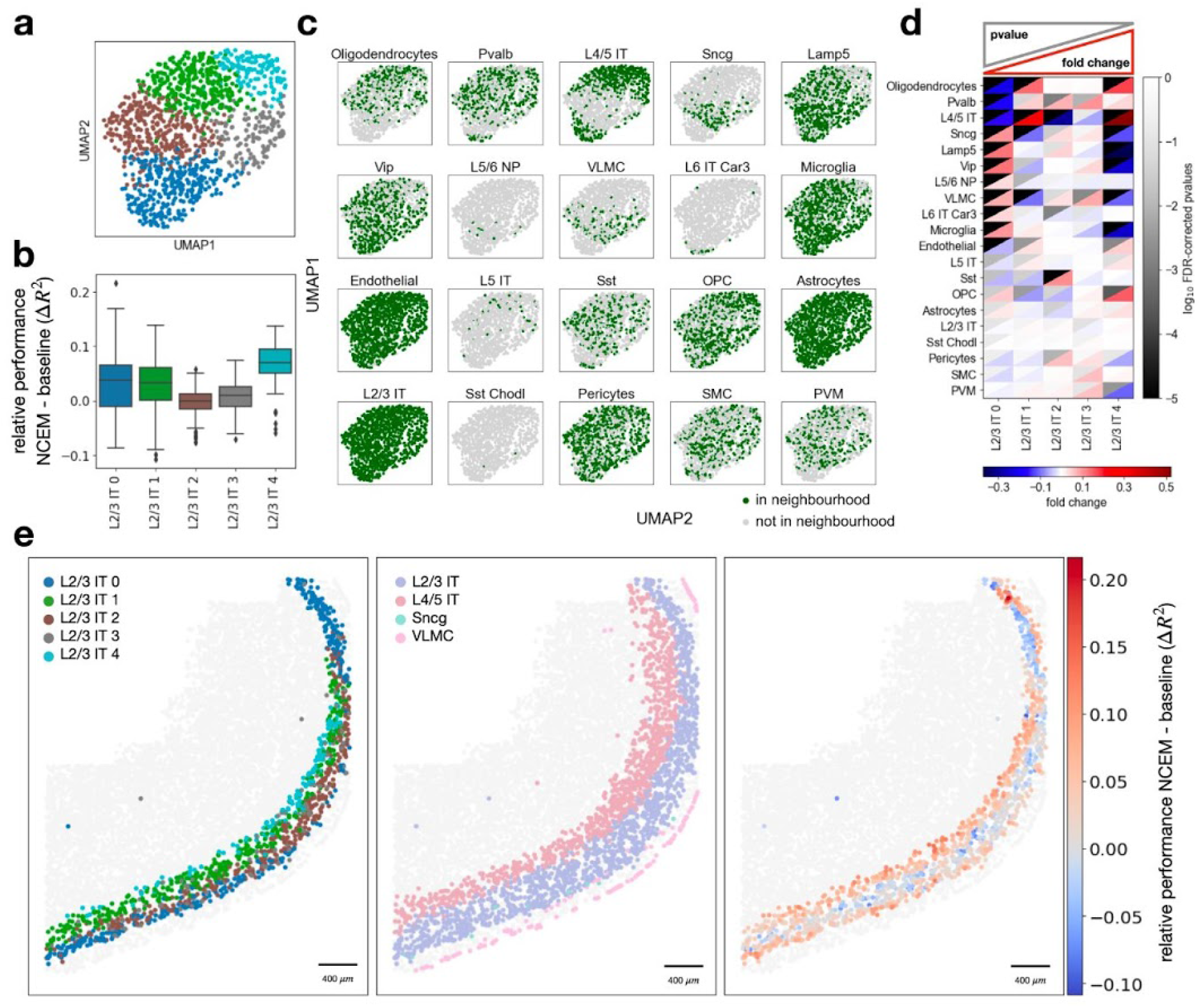
Cell heterogeneity can be attributed to niche composition. **(a)** UMAPs of molecular embedding of L2/3 IT cells with molecular sub-clustering superimposed (colors as in b). **(b)** Distribution of cell-wise difference of R^2^ between NCEM and non-spatial baseline model by molecular sub-cluster (L2/3 IT 0: n = 316, L2/3 IT 1: n = 314, L2/3 IT 2: n = 313, L2/3 IT 3: n = 133, L2/3 IT 4: n = 128). The centerline of the boxplots defines the mean over all relative R^2^ values, the height of the box is given by the interquartile range (IQR), the whiskers are given by 1.5 * IQR and outliers are given as points beyond the minimum or maximum whisker. **(c)** UMAPs of molecular embedding of all L2/3 IT cells in an example image (n = 1204 cells) showing if a given cell-type is present in the neighborhood. The underlying neighborhoods were defined at the optimal resolution defined in Fig. 1d (100 μm). **(d)** Heatmap of fold change versus false-discovery rate corrected p-values of cluster enrichment of binary neighborhood labels, where fold changes are the ratio between the relative neighboring source cell-type frequencies per subtype cluster and the overall source cell-type frequency in the image. **(e)** Spatial allocation of slice 153 of mouse brain in the MERFISH – brain dataset with L2/3 IT sub-states superimposed, L2/3 IT, L4/5 IT, Sncg, and VLMC superimposed and superimposed the difference of R^2^ between the NCEM at resolution of 100 μm and the best nonspatial baseline model.

In an equivalent analysis on the chip cytometry – colon data, we discovered a dependency of CD8 T cells on multiple other cell-types, such as CD8 T cells and lamina propria cells (sub cluster 3) (Supp. Fig. 8a-e), highlighting a dense compartment of T cells in the inflamed colon centred on lamina propria cells (Supp. Fig. 8f). In colorectal cancer, linear NCEM learned a previously established dependency of CD8 T cell states on proximity to the tumor–immune boundary^8^ (Supp. Fig. 9a-e), where sub cluster 1 represents cells close to malignant epithelial cells (Supp. Fig. 9c,f).

Next, we considered parameter significance in GLMs as a mechanism to attribute cellular variation to communication events with specific sender cell-types. We used a differential expression test, a Wald test, to test gene-wise cell communication coefficients in the linear model. In the linear model as defined above, the Wald test yields a p-value for the effect of the interaction between each pair of cell-types on each assayed gene in the receiver cell-type. The inference of cell communication with this gene-centric model improves with the number of assayed genes, as only a subset of all genes will display differential expression in response to the communication event. We found receiver cell-type-gene pairs that significantly depend on the neighborhood in the chip cytometry – colon data (Supp. Fig. 10c): Here, one would expect communication events with immune cells because of the inflamed state of the colon. The communication between sender PD1L+ cells and receiver CD8+ T cells was the strongest in terms of the number of differentially expressed genes, suggesting that PD1-mediated T cell exhaustion^20^ plays a role in this tissue. Importantly, this communication is not identifiable from cell contact frequencies (Supp. Fig. 6b) or CellphoneDB analysis (Fig. 7c), as expected for a sender–receiver-specific signature that occurs in a particular tissue niche. Indeed, we found PD1 in the set of genes whose expression is significantly associated with this receiver-sender pair (p=0.02 for an absolute change in log-transformed data of −1.92, Supp. Fig. 10).

### Nonlinear NCEM capture complex cell communication motifs

Linear models for cell communication cannot easily account for higher order interactions between more than two cell-types. Such complex interactions are, however, models of relevant patterns of communication in niches: The communication between two cells A and B may depend on a communication event between B and a cell C which induces ligand expression in B. Moreover, edge properties, such as weights derived from physical distances between cells, or molecular characteristics of communication axes, are difficult to account for in linear models. Here, we discuss multilayer NCEMs which can model both higher order cell communication and edge properties. These graph neural networks can be understood as encoder-decoder models. The encoder is a function of the local context in the graph and yields a latent state that is transformed by a stack of densely-connected layers in the decoder to the predicted expression state of the cell (Fig. 3a). In contrast to autoencoding encoder-decoder models often used to model heterogeneity in single cells from dissociated tissues^21,22^, this encoder-decoder NCEM is a nonlinear regression model that does not receive the expression state in the input (Online Methods).

**Figure 3:**
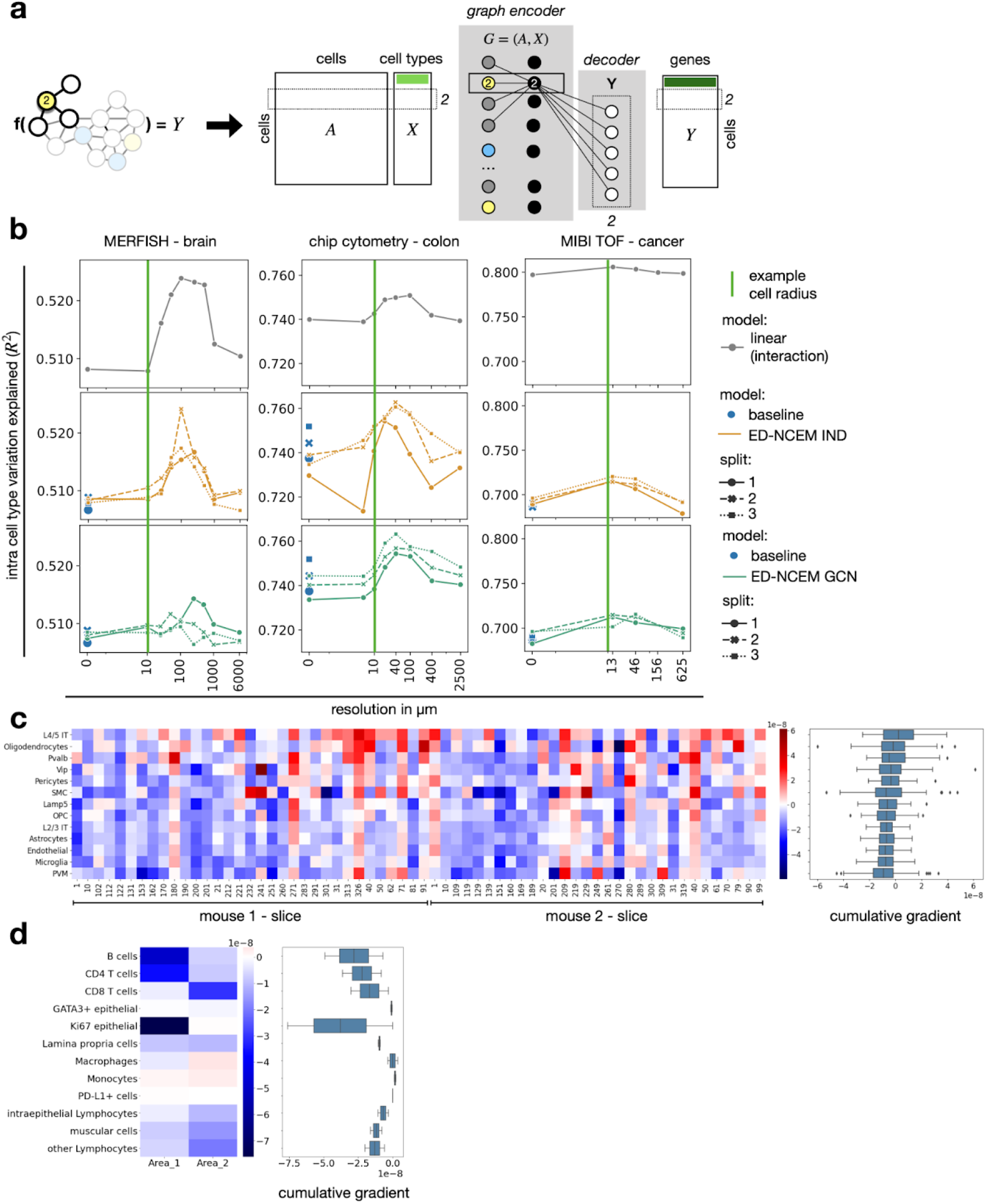
Nonlinear models of spatial dependencies of expression states. **(a)** A node-supervised model in which the supervision label is the expression vector of a cell and the input consists of categorical cell-type assignments and a spatial neighborhood graph. This model can also be viewed as an encoder-decoder model. The encoder performs a graph-based embedding of a single cell and the decoder translates a bottleneck activation vector into an expression state. **(b)** Nonlinear spatial dependencies in single-cell datasets. Shown are the R^2^ values for held-out test data of encoder-decoder models by resolution in μm with cross validation indicated as point shape and line style and comparatively mean performance of linear model in Fig. 1d. *Linear (interaction) (gray line)*: linear model with interaction effects; *ED*: encoder-decoder model; *IND*: the graph convolution is an indicator function across cell-types in the neighborhood (yellow lines); *GCN*: the graph convolution is a linear embedding (filter) of the cell-types in the neighborhood (teal lines); *split (point shapes)*:cross-validation split. **(c)** Heatmap of cumulative gradients (saliency) of gene expression prediction of L2/3 IT with respect to the input cells, aggregated by the sender cell-type clusters, on test data. Shown is a cumulative gradient matrix of L2/3 IT predictions by source cell-type and image. **(d)** Heatmap of cumulative gradients (saliency) of gene expression prediction of CD8 T cells with respect to the input cells, aggregated by the sender cell-type cluster, on the test data. Shown is a cumulative gradient matrix of CD8 T cell predictions by source cell-type and image. The cumulative absolute gradients in **(c, d)** are derived from the absolute gradients tensor across each cell’s molecular vector prediction with respect to the cells in the neighborhood (source cells) per image (tensor shape: *genes × cells × images*), by taking a sum across the molecular output features and by taking a sum across source cells of the same type (tensor shape: *cell – types × images*). For each box in **(c, d)**, the centerline defines the mean over all image-wise saliencies, the width of the box is given by the interquartile range (IQR), the whiskers are given by 1.5 * IQR and outliers are given as points beyond the minimum or maximum whisker.

First, we established the presence of resolution-dependent prediction performance profiles in encoder-decoder NCEMs in both models with indicator aggregators and graph-convolutional filters, again controlled by empty and large neighborhoods (Fig. 3b). The top-performing length scales were comparable to those obtained on linear models (Fig. 3b). We could identify cell-type-specific communication length scales again (Supp. Fig. 11). Notably, the encoder-decoder models did not outperform linear models on any dataset in terms of reconstruction metrics on test data, which suggests that niche communication is described well by additive pairwise communication events of cells in these tissues at the given sample complexity, rather than higher-order interactions.

As any neural network, encoder-decoder NCEMs can be interpreted in terms of cell communication events through gradient analysis. The gradients of output expression values with respect to input receiver cell-type and input sender cell-types approximate the effect that a change in neighborhood composition of a given cell would have on its molecular state (Online Methods). Using this interpretation method, we identified L4/5 IT as a putative sender cell-type for L2/3 IT cells in the MERFISH - brain dataset^12^ (Fig. 3c), which agrees with the strong association between these two cell-types identified in the cluster enrichment (Fig. 2c). Using a similar approach, we identified lamina propria cells as predictors of CD8 T cell state in the chip cytometry - colon dataset (Fig. 3d), reproducing this same association identified with linear NCEM (Supp. Fig. 8c,e).

### NCEMs extended to latent cell-intrinsic states explain more variance

Above, we discussed NCEMs as mechanistic models designed to attribute intra-cell-type variance to predictors in the niche. These NCEMs cannot model latent cell states that confound the measured expression data, representing important intrinsic cellular phenomena, such as cell cycle states or differentiation progression. A holistic model of cellular variance needs to account both for these intrinsic and extrinsic effects to disentangle confounded sources of variation^5^. Therefore, we propose an NCEM that accounts for both cell intrinsic latent states and the dependencies of molecular states on the niche. The proposed model is a conditional variational autoencoder (CVAE) in which the condition represents the neighborhood and the cell-type of the cell itself (Fig. 4a). In contrast to the node-supervised models discussed above, this CVAE is a node-generative model. This node-generation tasks of learning a distribution over gene expression states conditioned on the cell-type, a local context in the graph, and other covariates, such as chemical perturbation, extends generative models for single-cell gene expression data^21,23^ to spatial cell-cell dependencies.

**Figure 4:**
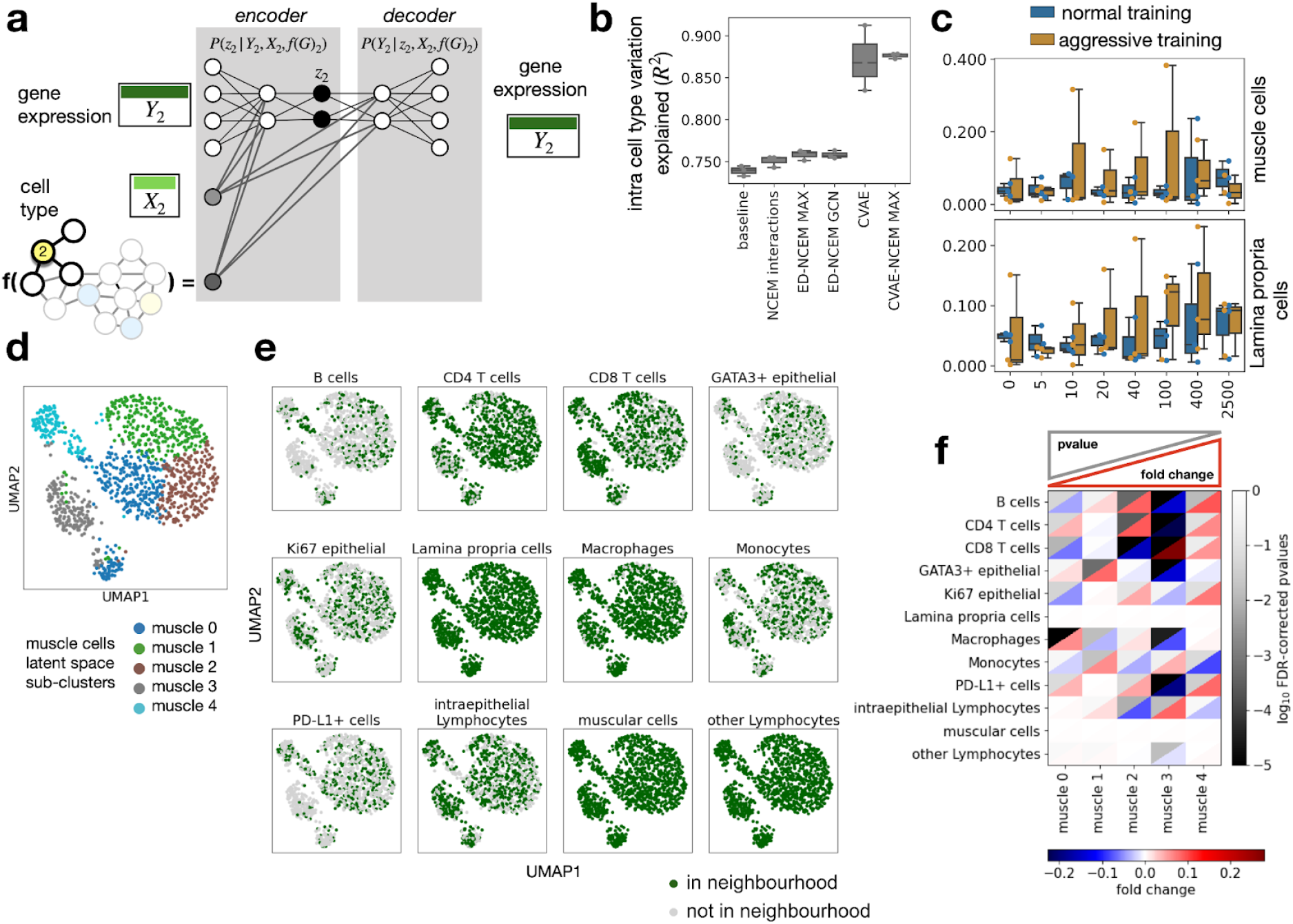
Modelling intrinsic and extrinsic variation in deep latent variable models. **(a)** A node generative network (CVAE–NCEM) is a conditional variational autoencoder in which the condition is not a constant but a graph embedding, which is also learned. **(b)** Latent variable models improve reconstructive performance. Shown are the R^2^ values of held-out test data based on the forward pass model evaluation from chip cytometry – colon data for linear models, encoder–decoder models, and variational autoencoders for both NCEM and nonspatial models. *baseline*: a nonspatial linear model of gene expression per cell-type; *NCEM interactions*: linear model with interaction effects; *ED*: encoder–decoder model; **IND**: the graph convolution is an indicator function across cell-types in the neighborhood; *GCN*: the graph convolution is a linear embedding (filter) of the cell-types in the neighborhood. **(c)** Neighborhood transfer performance of NCEM and nonspatial models. Shown is the R^2^ over cells in the test set for models trained on predicting muscular cells and Lamina propria cells for both CVAE and CVAE–NCEMs trained on neighborhoods with different radii with optimization algorithm as color. *Plain*: normal CVAE training; *aggressive*: aggressive encoder training. For each box in **(b, c)**, the centerline defines the mean over all three cross-validations, the height of the box is given by the interquartile range (IQR), the whiskers are given by 1.5 * IQR and outliers are given as points beyond the minimum or maximum whisker. **(d–f)** Latent variables of CVAE–NCEM are confounded with neighborhood conditions. **(d)** UMAP of molecular embedding in the CVAE–NCEM IND latent space of muscular cells in an example image (n = 1,149 cells) with molecular sub-clustering superimposed (muscle 0: n = 315, muscle 1: n = 287, muscle 2: n = 238, muscle 3: n = 183, muscle 4: n = 126)**. (e)** UMAPs of molecular embedding in the CVAE-NCEM IND latent space of all muscle cells in the same image with superimposed binary label of presence of a given cell-type, as defined in the title, in the neighborhood. The underlying neighborhoods were defined at a resolution of 100 μm. **(f)** Heatmap of fold change versus false-discovery corrected p-values of cluster enrichment of binary neighborhood labels where fold changes are the ratio between the relative neighboring source cell-type frequencies per subtype cluster and the overall source cell-type frequency in the image.

Conditional generative models have previously been used to model perturbations of molecular states by small molecules^23^ in the context of data from dissociated tissues. Latent variable models attain much higher predictive performance in reconstruction tasks (Fig. 4b, Supp. Fig. 13a) because they can fit cell-wise states if the model does not suffer from posterior collapse^24^. However, conditional variational models suffer from a non-identifiability between the variation attributed to the condition and to the latent states because the loss function only constrains the model’s marginal likelihood. A CVAE that is ideally suited for style transfer is converged such that the condition-wise marginal posteriors on the latent variables are equal, so that one can sample the full conditional distribution by decoding the latent states observed in a source condition into the target condition. However, there are optima of equal loss values in which the condition information bleeds into the latent variables, resulting in condition-specific latent states that cannot meaningfully be transferred to a target condition. We attempted to address this non-identifiability with aggressive encoder training, an optimization approach capable of favoring convergence to a model that attributes maximal variation in the marginal distribution to the conditional^25^ (Online Methods). We evaluated the ability of a CVAE-NCEM to extrapolate to unseen neighborhoods, a style-transfer setting we call “neighborhood transfer” (Online Methods) using the example of muscular and Lamina Propria cells from the chip cytometry – colon data (Fig. 4c) and L2/3 IT neurons from the MERFISH – brain dataset^12^ (Supp. Fig. 13b). We did not find consistent neighborhood-transfer performance differences between CVAE-NCEM compared and plain CVAEs, even though peak performance was attained by CVAE-NCEM optimized with aggressive encoder training (Online Methods). This performance analysis suggests that niche states are represented in latent variables, thus confounding resampling in the neighborhood transfer task. Indeed, we could identify multiple significantly enriched neighborhoods in a latent space clustering of the CVAE–NCEM for both examples (Fig. 4d-f, Supp. Fig. 13c-e). This latent variable interpretation shows that the CVAE indeed converges to an optimum that is not desirable in terms of the style transfer task, which can occur because these optimums are not penalized by the CVAE’s cost function.

NCEMs can attribute molecular variation of single-cells to niche composition and thus carry the promise of explaining variance previously not interpretable in dissociation data. Still, even with niche variance attributed, further sources of cellular heterogeneity are latent in spatial assays (Fig. 4b). CVAE–NCEMs are a model class that can close this gap between niche variation and latent variables. Further algorithmic work and targeted data acquisition are required to resolve the non-identifiability encountered in the examples discussed here.

## Discussion

We presented NCEMs, a graph neural network framework for modeling of cell communication events in spatial omics assays with subcellular resolution, in which cells can be separated *in silico* through segmentation. The NCEMs presented in this paper are structured in three groups: a linear graph neural network that can be mapped to a GLM, nonlinear graph neural network and CVAEs. We defined the extrapolation to unseen neighborhoods, the prediction of the molecular state of a cell of a given type in a given niche, as a supervision task and proposed an adaptation of this task to generative style transfer models (“neighborhood transfer”). We used NCEMs to discover statistical dependencies between cells on physiologically relevant length scales at a mean of 69 μm with a standard error of 14 μm across five datasets. We demonstrated that NCEMs could be interpreted based on model parameters in linear models and saliencies in nonlinear models to infer communication events between pairs of cell-types and to disentangle molecular variation in standard cell-centric unsupervised workflows. Using these interpretation mechanisms, we disentangled niche effects in the mouse motor cortex, in the inflamed colon and in colorectal cancer.

The datasets discussed here are based on technologies with high throughput in the number of cells but relatively few genes measured. For example, imaging-mass cytometry^26^ and CODEX^6^ both profile on the order of tens of genes. Typically, these genes are selected to separate cells from different cell-types, maximizing between-type variance in the data. However, these setups are not designed to profile within-type variance. The more subtle cell states within a cell-type are typically recovered in protocols with large feature spaces, such as spatial single-cell RNA-seq, but also FISH-based protocols, such as seq-FISH^27^ and MERFISH^9^. Although gene expression-based inference of cell communication will probably be improved by techniques that yield larger feature spaces, we could already show in this study that spatial dependencies can be estimated across cross-validation splits of the larger MERFISH dataset with 254 genes measured. We anticipate that such high-dimensional RNA-seq characterizations of single cells in tissues will soon be widely available because of the improvements of seq-FISH and MERFISH protocols and the reduction of spot sizes in spot-based transcriptomics. Datasets with more comprehensive quantification of ligand and receptor expression per cell may allow for further constraints on edges based on the expression of specific ligand and receptor pairs in the participating nodes. Such constraints may further increase interpretability and improve discovery of cell communication events through linear model interpretation and saliency-based analyses. Such constraints may further be derived through imputation of relevant ligand and receptor genes in spatial assays using reference single-cell atlasses^14^. All models proposed here are node-centric and embed a cell locally in the graph, instead of requiring the full graph. Therefore, NCEMs can be used equally well and efficiently on a few large images or on many small images, thereby increasing their relevance for a wider range of scenarios.

The datasets modeled here are 2D and do not contain information about cells that are also part of a receiver cell’s 3D neighborhood but are located in a different z-section; this limits the capacity of the models shown here to find communication events. One would expect NCEMs to be considerably stronger given 3D data, and we expect such data to become available from serial slicing of tissues, for example.

Here, we considered graph convolutional networks with linear and indicator graph aggregator functions. Other aggregators are used commonly to compute graph embeddings^28^ and could also be used here. The complexity of the graph neural network used in the NCEM defines the complexity of the motifs of cell communication that can be discovered. Linear NCEMs discover directional effects between pairs of cells. Higher order effects, that can be captured by deeper graph neural networks, include interactions between different communication events on a target cell and conditional dependencies between communication events, such as in loops. NCEMs do not require complex tissue phenotype labels, which are often unavailable. Effectively, the molecular vectors represent a label on a mid-range length scale of the graph. One can also imagine extending the tissue representation with more global phenotype labels through additional supervision tasks. The coarseness of the input feature space is an important hyperparameter for these graph models. Future work may address the choice of this hyperparameter algorithmically. However, in many cases, there is strong prior knowledge on the assignment of cell states to discrete cell-types.

The cell-cell dependencies modeled based on spatial graphs are examples for observational dependencies that violate the common assumption of statistical learning, namely, that single cells are independent and identically distributed. We showed that this assumption could be sacrificed given strong inductive priors, here induced by the spatial graphs, which constrains the modeled communication events from the set of all pairs of cells in a sample to the molecularly plausible ones. Future work could consider further inductive priors which are not necessarily spatial.

## Online Methods

### Data

#### MERFISH - brain

Zhang et al.^12^ measured mouse primary motor cortex with multiplexed error-robust fluorescence in situ hybridization (MERFISH) in 634 images across two mice with 254 genes observed in 284,098 cells. The cell-types originally annotated by Zhang et al., that are also used here, are astrocytes, endothelial, L2/3 intra-telencephalic (IT) neurons, L4/5 IT, L5/6 near-projecting (NP) neurons, L5 IT, L5 pyramidal tract (PT) neurons, L6 cortico-thalamic (CT) projection neurons, L6 IT, L6 IT Car3, L6b, Lamp5, microglia, oligodendrocyte precursor cells (OPC), oligodendrocytes, perivascular macrophages (PVM), pericytes, Parvalbumin (Pvalb), smooth muscle cells (SMC), Sncg, somatostatin (Sst), Sst Chodl, vascular leptomeningeal cells (VLMC), Vasoactive intestinal polypeptide (Vip), and other cells are annotated, where L identifies the layer (L1-L6) of the distinctive laminar structure based on cytoarchitectural features (Supp. Fig. 1a). Pvalb, Sst, Vip, Sncg and Lamp5 define five subclasses of GABAergic cells. We removed cells labeled as “*other*” from the dataset. The gene-wise mean-variance relationship does not indicate count noise (Supp. Fig. 13a). As domain information, we used an identifier for the respective mouse. The dataset has a lateral resolution of 109 nm per pixel. Zhang et al. used a seeded watershed algorithm to identify cell segmentation boundaries in each image. They performed graph-based Louvain community detection^19^ with the first 35 principal components using Scanpy^17^ for k = 10 neighbors for cell-type clustering.

#### chip cytometry - colon

Jarosch et al. measured an inflamed colon with chip cytometry in two images across one patient with 19 genes observed in 11,321 cells. The cell-types originally annotated by Jarosch et al., which are modeled here, are B cells, CD4 T cells, CD8 T cells, GATA3+ epithelial, Ki67 high epithelial, Ki67 low epithelial, lamina propria cells, macrophages, monocytes, PD-L1+ cells, intraepithelial lymphocytes, muscular cells and other lymphocytes are annotated (Supp. Fig. 1a). We coarsened the cell-type annotation by combining Ki67 high epithelial and Ki67 low epithelial to a joined annotation of Ki67 epithelial. The gene-wise mean-variance relationship indicates count noise (Supp. Fig. 13b). Therefore, we log-transformed gene expression values. Jarosch et al. performed thresholding, watershed algorithm and additionally implemented a cell-type-specific segmentation method to segment individual cells. Intensity values were corrected for spatial spillover prior to quantification. Cell-types were clustered using the Leiden clustering^18^ of the neighborhood-graph.

#### MIBI TOF - cancer

Hartmann et al.^8^ measured colorectal carcinoma and healthy adjacent tissue with multiplexed ion beam imaging by time-of-flight (MIBI-TOF) in 58 images across four individuals with 36 genes observed in 63,747 cells. The cell-types originally annotated by Hartmann et al., which are modeled here, are endothelial, epithelial, fibroblast, CD11c myeloid, CD68 myeloid, CD4 T cells, CD8 T cells and other immune cells are annotated (Supp. Fig. 1a). A coarser cell-type labeling was not applied to this dataset. The cohort in this dataset includes two patients with colorectal carcinoma and two healthy controls. The images have a size of 400 μm^2^ and 1,024 × 1,024 pixels. The gene-wise mean-variance relationship does not indicate count noise (Supp. Fig. 13b). We scaled the model outputs by cell-wise size factors. Hartmann et al. trained a convolutional neural network. They fed the output into the watershed algorithm to segment individual cells and cell-types were clustered using the FlowSOM R package and manually annotated based on their lineage marker profiles.

#### MELC - tonsils

Pascual-Reguant et al.^7^ measured tonsils from patients undergoing tonsillectomy with multi-epitope ligand cartography (MELC), an immunohistochemistry approach, in one image across one patient, with 51 genes observed in 9,512 cells. The cell-types originally annotated by Pascual-Reguan et al., which are modeled here, are B cells, endothelial cells, ILC, monocytes/macrophages/DC, NK cells, plasma cells, T cytotoxic cells, T helper cells and other cells are annotated (Supp. Fig. 1a). We removed cells labeled as “*other*” from the dataset. The gene-wise mean-variance relationship does not indicate count noise (Supp. Fig. 13d). Pascual-Reguant et al. performed segmentation by applying a signal-classification step using Ilastik 1.3.2 and an object-recognition step using CellProfiler 3.1.8, which were analyzed and clustered in Orange 3.26.0.

#### CODEX – cancer

Schürch et al.^13^ measured advanced-stage colorectal cancer with co-detection by indexing (CODEX) in 140 images across 35 patients with 57 genes observed in 272,266 cells. The cell-types originally annotated by Zhang et al., which are modeled here, are B cells, CD11b+ monocytes, CD11c+ dendritic cells, CD11b+CD68+ macrophages, CD163+ macrophages, CD68+ macrophages, CD68+ macrophages GzmB+, CD68+CD163+ macrophages, CD3+ T cells, CD4+ T cells, CD4+ T cells CD45RO+, CD4+ T cells GATA3+, CD8+ T cells, NK cells, T regs, adipocytes, dirt, granulocytes, immune cells, immune cells / vasculature, lymphatics, nerves, plasma cells, smooth muscle, stroma, tumor cells, tumor cells / immune cells and vasculature are annotated (Supp. Fig. 1a). Cells with an annotation of dirt or an undefined label were removed from the dataset. A coarser cell-type grouping was applied to the macrophage groups CD11b+CD68+ macrophages, CD163+ macrophages, CD68+ macrophages, CD68+ macrophages GzmB+ and CD68+CD163+ macrophages. Additionally, CD4+ T cells, CD4+ T cells CD45RO+ and CD4+ T cells GATA3+ were grouped into CD4+ T cells. The gene-wise mean-variance relationship indicates count noise (Supp. Fig. 13e). Therefore, we scaled model outputs by the node size in the respective output layer of each model class. Schürch et al. performed segmentation using the CODEX toolkit segmenter and unsupervised cell-type clustering using X-shift.

**Table 1:** Overview of datasets analyzed in this study. Shown are the spatial molecular profiling chemistry, the domain effect accounted for via batch covariates, the data transform used on the expression vectors, the inclusion of cell size factors, the number of images given to the models during each update (batch size) and the number of nodes evaluated per image per batch (n).

| dataset /<br>first author | technology | domain | transform | node size<br>scaling | batch size | n |
| --- | --- | --- | --- | --- | --- | --- |
| Zhang | MERFISH | patient | - | False | 64 | 10 |
| Jarosch | Chip Cytometry | patient | log(x+1) | False | 2 | 100 |
| Hartmann | MIBI-TOF | image | - | True | 58 | 10 |
| Pascual-Reguant | MELC | patient | - | True | 1 | 200 |
| Schürch | CODEX | patient | log(x+1) | True | 140 | 10 |

### Models

The inputs to NCEMs are a gene expression matrix *Y* ∈ *R^N×J^*, where *N* is the number of cells and *J* is the number of genes with *y_i_* being the gene expression vector for gene *i* = 1,…, *J*, a matrix of observed cell-types *X_l_* ∈ *R^N×L^* where *L* is the number of distinct cell-type labels and *X_c_* ∈ *R^N×C^* is a matrix specifying the batch assignments, with *C* being the number of distinct batches or domains, such as images or patients, in the dataset. We denote the adjacency matrix of connected cells as *A* ∈ *R^N×N^* which is calculated based on the spatial proximity of cells per image. For linear models and models with an indicator aggregator, a binary adjacency matrix is used with *a_ij_* = 1 if *d*(*x_i_*, *x_j_*) ≤ *δ_max_* where *d*(·,·) describes the euclidean distance between nodes *i, j* ∈ *N* and *δ_max_* is the maximal distance between interacting nodes (neighborhood size), and *a_ij_* = 0 otherwise. For models using a GCN as graph layer, we normalize *A* such that all rows sum to one, so *D*^−1^ *A* where *D* is the diagonal node degree matrix. The output of NCEMs is 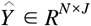, a reconstruction of the input count matrix *Y*, with 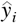 being the reconstructed expression for gene *i* = 1,…, *J*. For MIBI-TOF - cancer, MELC - tonsils and CODEX - cancer, we applied size factor scaling to the network output 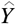. Let 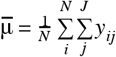 be the global mean per node, then size factors are given by 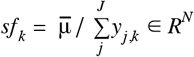. The network output scaling is then given by 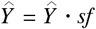.

#### Loss functions

According to the noise structure of the datasets explored in this paper (Supp. Fig. 13) we use a Gaussian log-likelihood loss as an optimization objective for GLM and ED models with 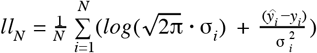, where σ_*i*_ is the predicted standard deviation for a gene *i* = 1,…,*N*. The loss function of CVAE models is the negative log-likelihood regularized by the Kullback-Leibler divergence between the encoder distribution *q*_φ_(*z*|*Y*) and the distribution of the latent space *p*(*z*), so *l_CVAE_ = − ll_N_* + *KL*[*q*_φ_(*z*|*Y*) ∥ *p*(*z*)]. For chip cytometry - colon and CODEX - cancer we therefore *log*(*Y* + 1) transformed the gene expression data.

#### Optimization

For each dataset, we ran grid searches to find the optimal set of hyperparameters. As batch size we choose the number of images per dataset. The number of nodes evaluated per image per batch was selected to ensure convergence and stabilize training. All models were trained with the Adam optimizer algorithm implemented in tensorflow. Linear models were trained with a *lr* = 0.5, the remaining models were trained for a varying learning rate of *lr* = {0.5, 0.05, 0.005}. Additionally, we used a learning scheduler on the validation loss with a patience of 20 epochs which reduces the learning rate by a factor of 0.5, so *lr_new_* = *lr* * 0.5 and early stopping with a patience of 100 epochs. The exact description of all grid searches in code are supplied in the benchmarking repository (Code Availability).

#### Linear NCEM

The NCEM includes two linear regression models. The nonspatial baseline linear model infers a reconstruction 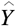 from a nodes cell-type and respective domain information via 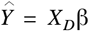, where *X_D_* is the design matrix and β ∈ *R*^(*L+C*)×*N*^ are the parameters learned by the model. The design matrix for nonspatial baseline models is given by *X_D_* = (*X_l_, X_c_*) ∈ *R*^*N*×(*L+C*)^. The spatial counterpart model, the NCEM, has access to an interaction matrix. First, we compute discrete target cell interactions with *X_T_* = 1_(*A*·*X_l_* > 0)_ ∈ [0, 1]^*N×L*^, where 1_(·)_ represents an indicator function. To generate a matrix representation of target-source cell interactions, we compute the interaction between each column of *X_l_* and each column *X_T_* via the point-wise product.The resulting interaction matrix is then denoted as 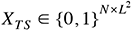 and the design matrix for the linear model with interaction terms is given by 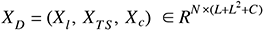. Equivalently, the model infers 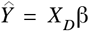 where 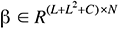 and additionally the gene variance per node which is inferred in the last layer of the linear model. We also considered an NCEM without interaction terms which does not have receiver-specific sender effects, but only global sender effects which account for the presence of senders in the niche via *X_S_* ∈ {0,1}^*N×L*^ : *X_D_* = (*X_l_*, *X_S_*, *X_c_*) ∈ *R*^*N*×(*L+L+C*)^. Coefficient significance is computed with Wald hypothesis testing with a significance threshold of τ = 0.01 on the parameters learned by the model for the interaction matrix *X_TS_*.

#### Nonlinear encoder-decoder NCEM (ED-NCEM)

The NCEM includes nonlinear encoder-decoder models that encode the neighborhood through a graph neural network (ED-NCEM) and decode expression vectors. The nonspatial null model is a nonlinear model (ED) that predicts expression from cell-type and graph-level predictors, alone. An encoder is given by *f_enc_* : *q*_φ_(*z* | *X_l_*, *g*(*A,X_l_*), *X_c_*), which encodes the cell-type labels *X_l_*, some graph-level predictors *X_c_* and the local graph embedding *g*(*A,X_l_*), based on the adjacency matrix *A*, into a latent state *z*. The latent state is input to the decoder and given by 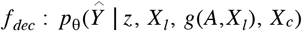. If *g*(*A,X_l_*) is and indicator embedding function as described in the section “Linear NCEM”, then *g*(*A,X_l_*) = *X_TS_* and the input to the linear model and the encoder are the same. If then also all hidden layers are removed from the ED-NCEM, a single linear transformation of the input remains which is equivalent to the linear NCEM. Alternatively, *g*(*A, X_l_*) can be a graph embedding learned by a graph-convolutional network (GCN)^10,11^. A one-layer GCN is given by 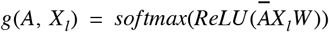 where *W* ∈ *R^L×H^* is an input-to-graph-embedding weight matrix with *H* being the dimension of the learned graph representation and 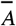 being the normalized adjacency matrix. In this case, *W* can be learned using gradient descent.

#### Conditional variational autoencoder NCEM (CVAE-NCEM)

A variational autoencoder^29^ learns a distribution over node states *Y* through a variational posterior over a latent space representation *z* which yields a reconstruction 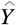 of *Y* via a likelihood function (the decoder). The nonspatial CVAE null model contains the cell-type and graph-level predictors as a condition in the variational posterior and the likelihood. In CVAE-NCEM, the conditions are the cell-type labels *X_l_*, some graph-level predictors *X_c_* and the local graph embedding *g*(*A,X_l_*), based on the adjacency matrix *A*. The encoder is then given by 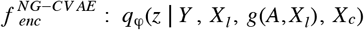 and the decoder is defined by 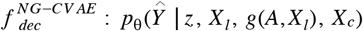. A CVAE–NCEM for a full dataset depends on both the spatial context and the type of the cell itself. This setting presents the challenge of encountering a non-identifiability between variance attributed to latent variables, cell-type conditions, and neighborhood context. In this study, we consider the CVAE–NCEM trained on the molecular vectors of a single target cell-type as a function of the full neighborhood context to remove the non-identifiability with respect to cell-type variation and focus on the non-identifiability between latent variables and neighborhoods.

### Normalized saliency maps

Saliency maps are used to differentiate the importance of features in the network input to analyze their importance for the network output for nonlinear models. In our case, saliency maps are aggregated at the cell-type level to extract communication events learned by the model, so *SALS* ∈ *R*^*L×L*^ with *L* being the number of distinct cell-types in the model. Non-normalized saliencies will show a pattern similar to the contact frequency matrix *M* ∈ *R*^*L×L*^ as cell-types with frequent connections will skew the learned importance of cell connections. We therefore normalize the saliencies by the absolute frequency of cell–cell connections, that is 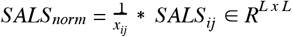 for *x_ij_* ∈ *M* with *i, j* = 1, …, *L*.

### Model evaluation

Overall model performance evaluation is based on the coefficient of determination 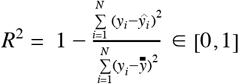 with 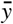 being the mean over gene expression values. Best performing models are selected based on highest *R*^2^ for the validation dataset. These metrics are based on model predictions, which derive from a forward pass through the model. The performance of CVAEs is additionally assessed in style transfer tasks. In style transfer, the gene expression state and neighborhood of a reference node from the source domain is encoded to estimate the latent states of this node. This latent representation is then decoded to the target domain, which implies conditioning the decoding on the target neighborhood:

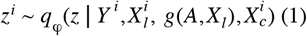

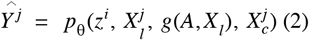

where *i, j* are cell indices, *q*_φ_ is the variational posterior and *p*_θ_ is the decoder network. See also “*Conditional variational autoencoder NCEM (CVAE–NCEM)*” for details on the notation.

### Ligand–receptor association analysis

For ligand–receptor permutation tests we used the CellphoneDB^3^ implementation in Squidpy^15^. For the chip cytometry – colon, MIBI TOF – cancer^8^, MELC – tonsils^7^, and CODEX – cancer^13^ datasets, node feature names are mapped to HGNC gene names. After the mapping, we used the ligand–receptor interaction pairs of the Omnipath database^30^ in Squidpy^21^ and ran the permutation test for n = 1,000 permutations. For three datasets, we used a random subsample of all cells (MERFISH – brain^12^ 10% with n = 27,655, MIBI TOF – cancer^8^ 40% with n = 25,498 and CODEX – cancer^13^ 10% with n = 25,186). Results are visualized with Squidpy^21^ with only p-values below a threshold of 0.3 shown.

### Variance attribution analysis

We used Uniform Manifold Approximation and Projection^31^ (UMAP) to embed the cells in two dimensions for visualization of high-dimensional data. For the UMAPs of the MERFISH – brain data^12^ matrix (Fig. 1c and Supp. Fig. 5b) we performed dimensionality reduction using PCA with the first 35 principal components (PCs) and the nearest neighborhood size of k = 10. A similar approach was described by Zhang et al. to identify stable clusters for subsequent cell-type annotation. For the UMAPs of L2/3 IT neurons in slice 153 (Fig. 2) and slice 162 (Supp. Fig 5b) of the MERFISH - brain dataset, we used the first 40 PCs with k = 40 and performed graph-based Louvain community detection^19^ using Scanpy^17^ to define stable L2/3 IT substates. For the UMAPs of CD8 T cells in area 1 in the chip cytometry dataset (Supp. Fig. 8), we used the input matrix directly and k = 22. For the

UMAPs of CD8 T cells in image 1, 5, 8 and 16 of the MIBI-TOF - cancer dataset (Supp. Fig. 9), we used the input matrix directly and k=60. Clustering of the latent space in CVAE and CVAE-NCEM IND models (Fig. 4d, Supp. Fig. 11c) was performed using the latent space matrix directly and k = 80 for the MERFISH - brain dataset and k = 250 for the chip cytometry - colon dataset.

For cluster enrichment analysis, we performed Fisher’s exact test. Each contingency table is composed of two categorical variables. The first variable is the number of cells in one specific L2/3 IT substate versus the remaining L2/3 IT substates. The second variable is the number of cells with a respective source type in their neighborhood and those cells where this source type is not present in the neighborhood. We performed Benjamini and Hochberg false discovery rate correction (FDR) of cluster enrichment p-values. A similar approach was used for the cluster enrichment analysis of CD8 T cells in the chip cytometry - colon and the MIBI-TOF - cancer datasets.

For comparison of the source cell-type rankings of contact frequencies, ligand-receptor analysis and cluster enrichment analysis of L2/3 IT cells, we performed Kendall’s tau correlation analysis and computed the tau statistic and the two-sided p-value. Contact frequencies are ranked by source type frequencies (row L2/3 IT of Supp. Fig. 6). For ligand-receptor analyses, source cell-types are ranked based on the number of significant p-values below a threshold of 0.05 for this L2/3 IT interaction. For the cluster enrichment analysis we used the ranking shown in Fig. 2d.

#### Variance decomposition into inter- and intra-cell-type variance

The variance of a single-cell resolved dataset can be decomposed into inter-cell-type variance, intra-cell-type variance, and gene variance. The gene variance is independent of cell-type definitions and can therefore be considered separately from relative intra- and inter-cell-type variance.

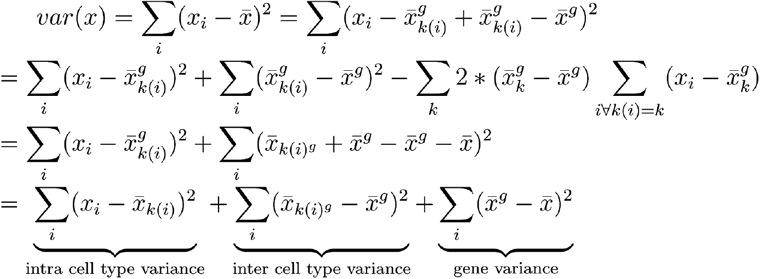

where x_i_ is the expression vector of cell *i*, 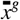 is the vector of gene-wise means of the dataset, 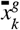 is the vector of gene-wise means of the cells in cell-type *k*, *k(i)* is the cell-type of cell *i*, and 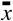 is the scalar gene- and cell-wise mean of the dataset.

## Data Availability

The MERFISH - brain^12^, MIBI TOF - cancer^8^, MELC - tonsils^7^ and CODEX - cancer^13^ datasets are publicly available (Online Methods). The chip cytometry - colon dataset has been generated by the Busch lab and is currently under review, and has kindly been provided to us.

## Code Availability

All models described here are implemented in a Python package available at https://github.com/theislab/ncem. All benchmarking and analysis codes are provided at https://github.com/theislab/ncem_benchmarks. Tutorials for model usage are available from https://github.com/theislab/ncem_tutorials.

## Author Contributions

DSF and FJT conceived the project. DSF and ACS performed the analysis and wrote the code. DSF, ACS and FJT wrote the manuscript.

## Acknowledgements

We would like to thank Sabrina Richter, Mohammad Lotfollahi, Prof. Dr. Stephan Günnemann, Dr. Christian M. Schürch, Sebastian Jarosch and Prof. Dr. Dirk Busch for valuable discussion and feedback on this project. In particular, we want to thank Sebastian Jarosch and Prof. Dr. Dirk Busch for sharing the chip cytometry – colon dataset pre-publication. We would like to thank Dr. Luke Zappia and Giovanni Palla for their valuable feedback on this manuscript.

This work was supported by the German Federal Ministry of Education and Research (BMBF) under Grant No. 01IS18036B and No. 01IS18053A, by the Bavarian Ministry of Science and the Arts in the framework of the Bavarian Research Association “ForInter” (Interaction of human brain cells), by the Wellcome Trust Grant 108413/A/15/D and by the Helmholtz Association’s Initiative and Networking Fund through Helmholtz AI [grant number: ZT-I-PF-5-01]. D.S.F. acknowledges support from a German Research Foundation (DFG) fellowship through the Graduate School of Quantitative Biosciences Munich (QBM) [GSC 1006 to D.S.F.] and by the Joachim Herz Foundation. ACS has been funded by the German Federal Ministry of Education and Research (BMBF) under Grant No. 01IS18036B.

