## Supplemental Figures for "Learning cell communication from spatial graphs of cells"

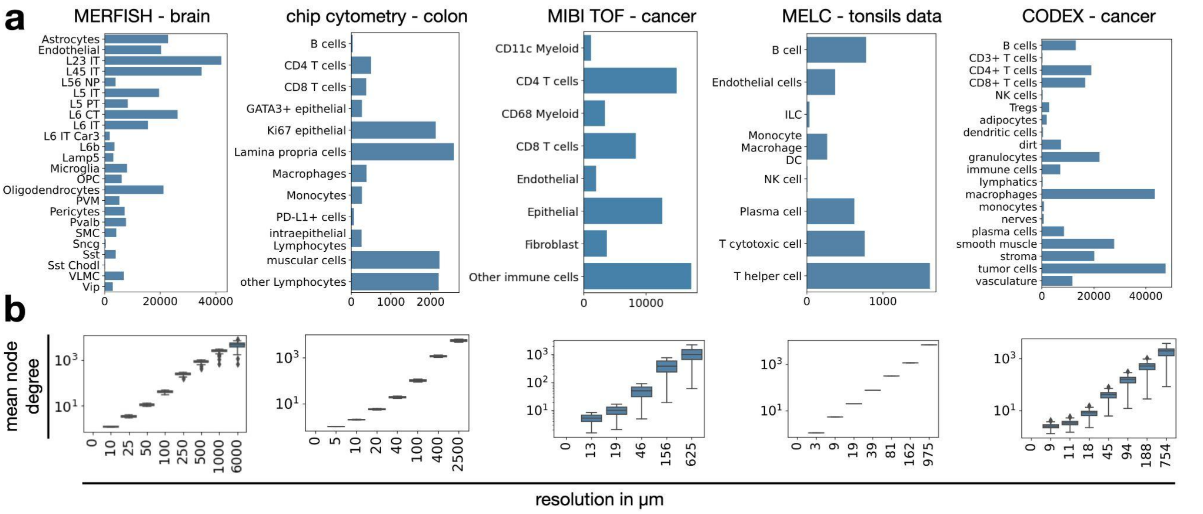


**Supp. Fig. 1: Cell-type centric summary statistics per dataset. (a)** Cell-type frequencies by dataset. Shown is a barplot with the number of cells in each cell-type for MERFISH – brain data, chip cytometry – colon data, MIBI TOF – cancer data, MELC – tonsils data and CODEX – cancer data. **(b)** Mean node degree (number of neighbours) by resolution in µm and dataset for MERFISH – brain data, chip cytometry – colon data, MIBI TOF – cancer data, MELC – tonsils data and CODEX – cancer data. For each box in **(b)**, the centerline defines the mean over all images the height of the box is given by the interquartile range (IQR), the whiskers are given by 1.5 * IQR and outliers are given as points beyond the minimum or maximum whisker.

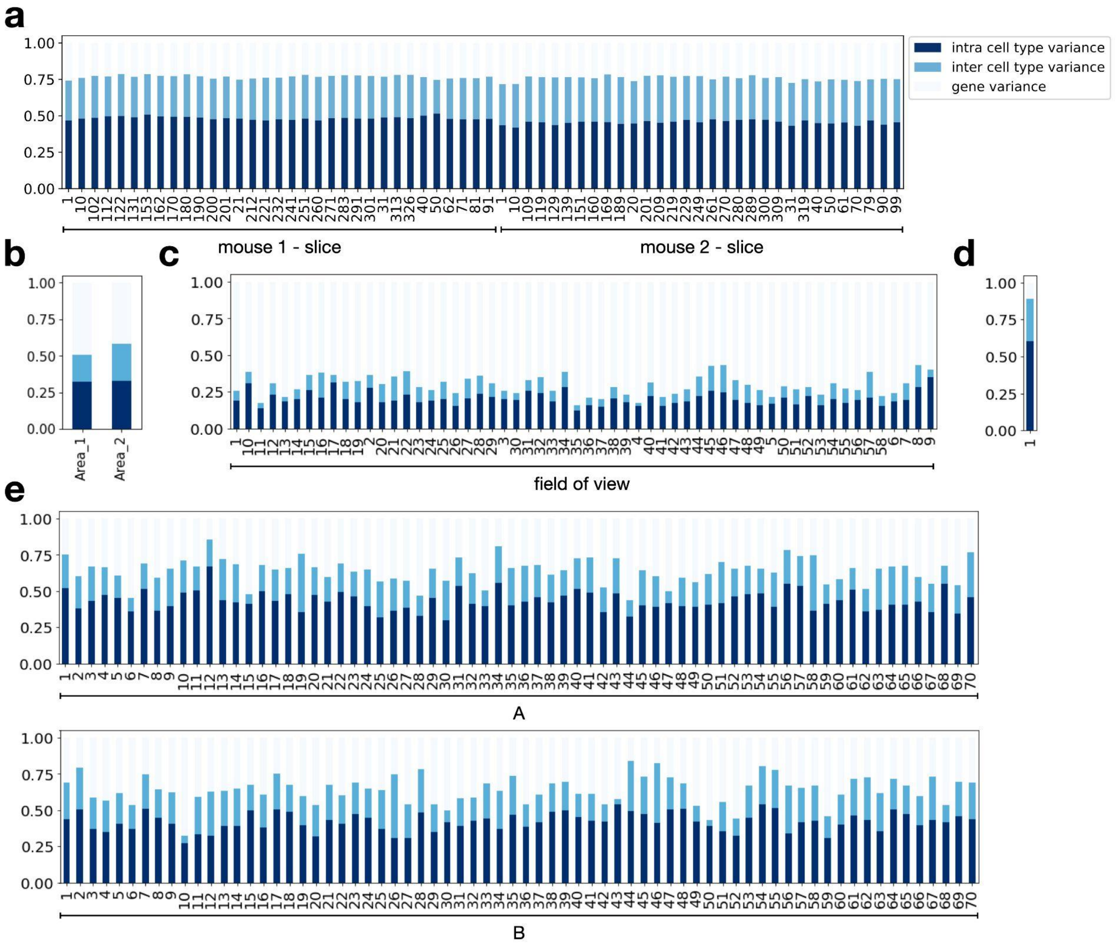

**Supp. Fig. 2: Variance decomposition in spatial omics datasets.** MERFISH – brain (mean intra cell type variance: 47%, mean inter cell type variance: 29%, mean gene variance: 24%) **(a)**, chip cytometry – colon (mean intra cell type variance: 33%, mean inter cell type variance: 22%, mean gene variance: 45%) **(b)**, MIBI TOF – cancer (mean intra cell type variance: 20%, mean inter cell type variance: 10%, mean gene variance: 70%) **(c)**, MELC – tonsils (mean intra cell type variance: 60%, mean inter cell type variance: 29%, mean gene variance: 11%) **(d)** and CODEX – cancer dataset (mean intra cell type variance: 43%, mean inter cell type variance: 21%, mean gene variance: 36%) with images ordered by tissue microarrays (*A, B*) **(e)**.

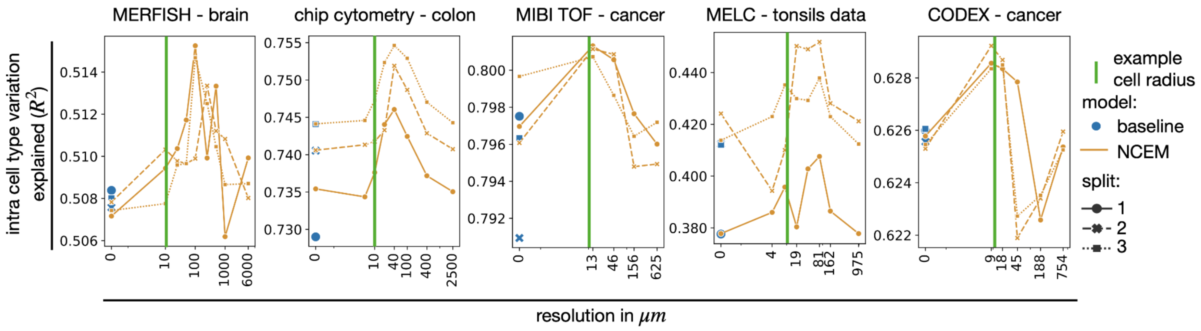


**Supp. Fig. 3: Linear models for spatial cell state dependencies.** Linear models without receiver-sender interaction terms (Online Methods) capture neighborhood dependencies in spatially resolved single-cell data. Shown are R2 for held-out test data of linear models by resolution in µm with cross validation indicated as point shape and line style. The underlying linear models are parameterised with sender cell-type-specific parameters. *example cell radius (green line)*: Example length scale of a cell, here chosen as 10 µm; *baseline (blue dot)*: a nonspatial linear model of gene expression per cell-type; *NCEM*: linear NCEM.

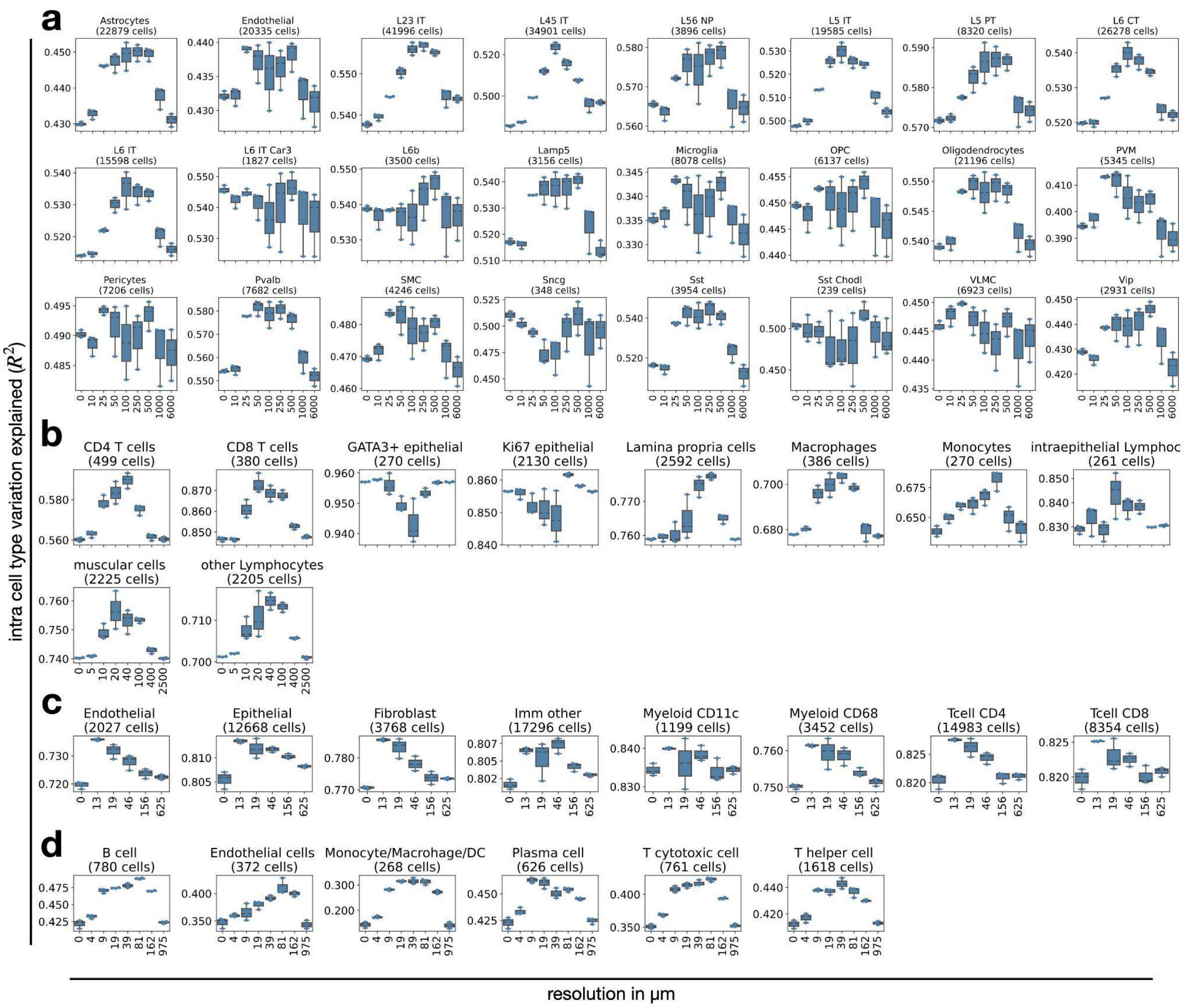


**Supp. Fig. 4: Length scales of dependencies for different target cell-types.** Shown are $R^2$ for held-out test data of linear models by resolution in µm for different predicted cell-types for MERFISH – brain data **(a)**, chip cytometry – colon data **(b)**, MIBI TOF – cancer **(c),** MELC – tonsils **(d)** where each boxplot corresponds to a three-fold cross validation. For each box in **(b-d)**, the centerline defines the mean over all three cross validations, the height of the box is given by the interquartile range (IQR), the whiskers are given by 1.5 * IQR and outliers are given as points beyond the minimum or maximum whisker.

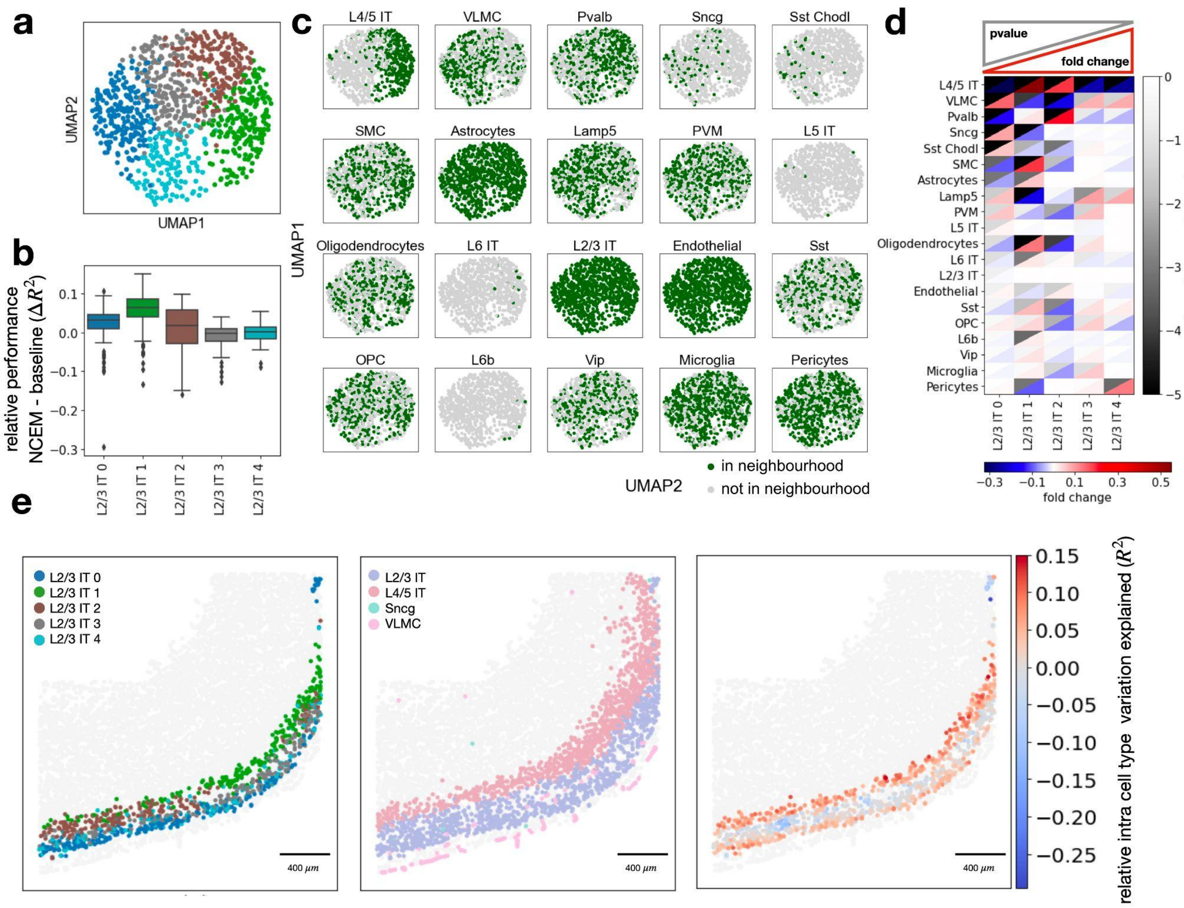


**Supp. Fig. 5: Attribution of molecular states to neighborhoods in the MERFISH – brain dataset. (a-e)** Replicate analysis corresponding to results presented in Fig. 2 on a second image from the MERFISH – brain dataset. **(a)** UMAPs of molecular embedding of L2/3 IT cells only with molecular sub-clustering superimposed (colors as in b)**. (b)** Distribution of cell-wise difference of $R^2$ between spatial model non non-spatial baseline model by molecular sub-cluster (L2/3 IT 0: n = 226, L2/3 IT 1: n = 209, L2/3 IT 2: n = 193, L2/3 IT 3: n = 191, L2/3 IT 4: n = 127). The centerline of the boxplots defines the mean over all relative $R^2$ values, the height of the box is given by the interquartile range (IQR), the whiskers are given by 1.5 * IQR and outliers are given as points beyond the minimum or maximum whisker. **(c)** UMAPs of molecular embedding of all L2/3 IT cells in example image (n = 946 cells) showing if a given cell-type is present in the neighborhood. The underlying neighborhoods were defined at the optimal resolution identified in Fig. 1d (100 µm). **(d)** Heatmap of fold change versus false-discovery rate corrected p-values of cluster enrichment of binary neighborhood labels where fold changes are the ratio between the relative neighboring source cell-type frequencies per subtype cluster and the overall source cell-type frequency in the image. **(e)** Spatial allocation of slice 162 of mouse brain in the MERFISH – brain dataset with L2/3 IT sub-states superimposed, L2/3 IT, L4/5 IT, Sncg, and VLMC superimposed and superimposed the difference of $R^2$ between the NCEM interaction model at a resolution of 100 µm and the best nonspatial baseline model.

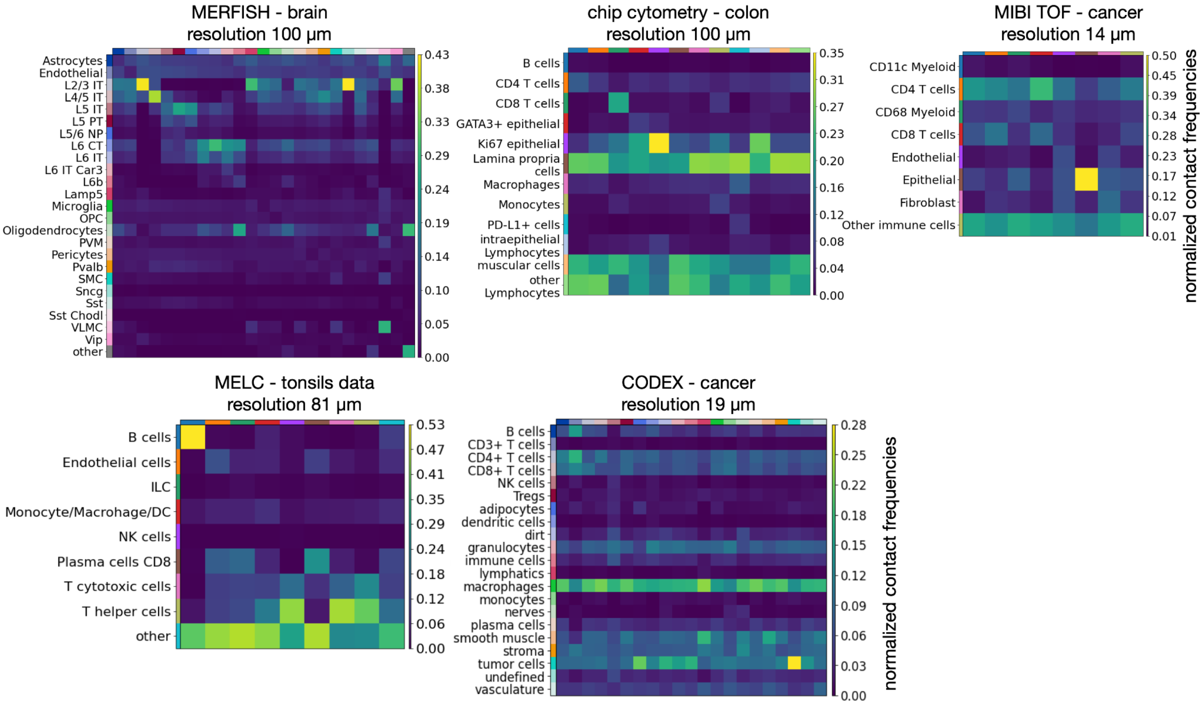


**Supp. Fig. 6: Normalised contact frequencies of cell-types in spatial neighborhoods.** The underlying resolution in µm was chosen based on the optimal linear NCEM performance (Fig. 1d). Normalised contact frequencies per data set are calculated as the mean of the image-wise normalised sender-receiver cell interaction extracted with Squidpy[15].

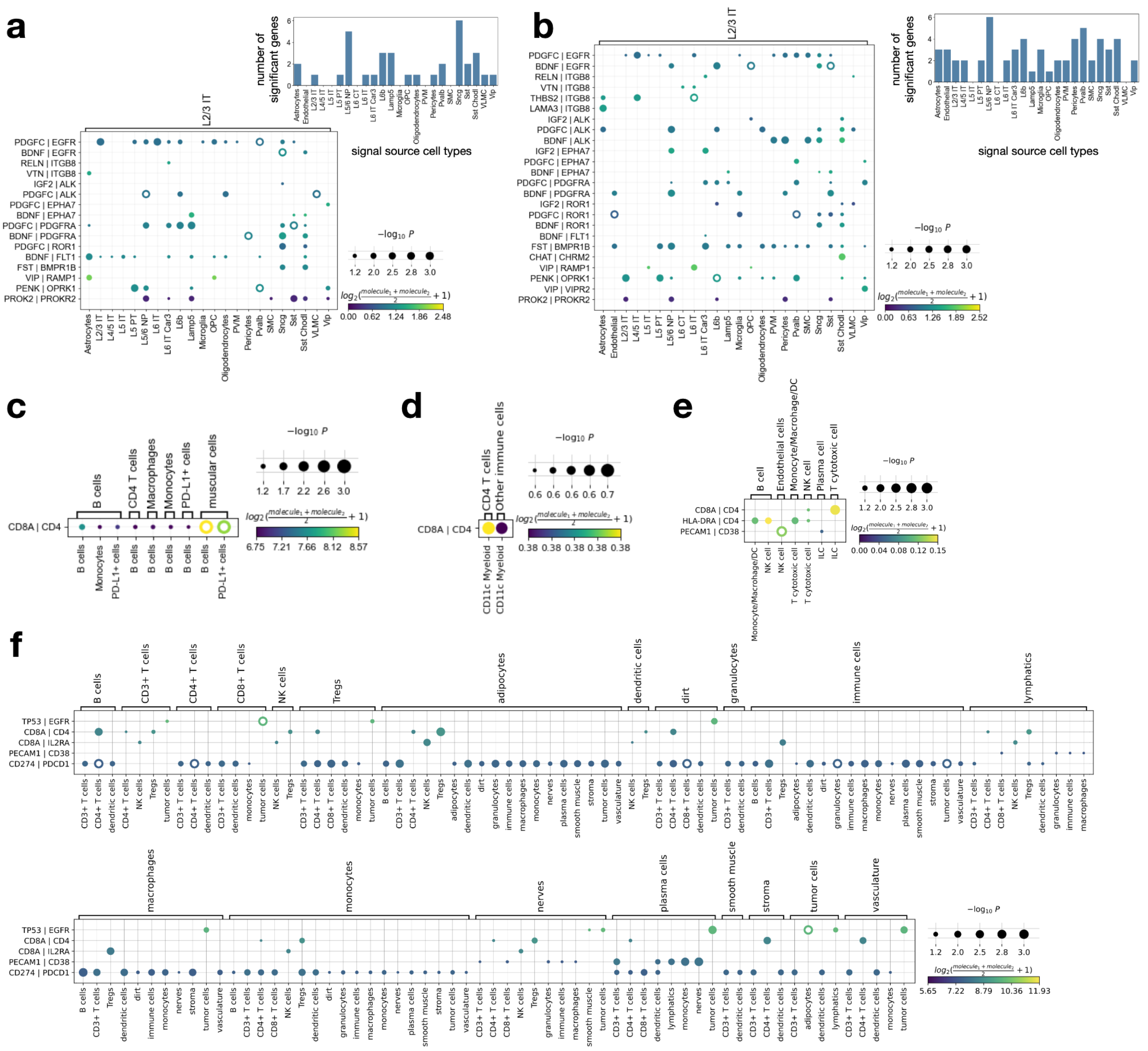


**Supp. Fig. 7: Ligand–receptor permutation test.** Shown are ligand–receptor gene pairs observed in the study and their significant associations across cell-types[3] for pairs with a p-value ≤ 0.3 for MERFISH – brain (complete dataset L2/3 IT neurons) **(a)**, MERFISH – brain (slice 153 L2/3 IT neurons) **(b)**, chip cytometry – colon **(c)**, MIBI TOF – cancer **(d)**, MELC – tonsils **(e)**, and CODEX – cancer **(f)**.. The barplots in **(a, b)** correspond to the count over non-corrected p-values below a threshold of 0.05 per L2/3 IT-source pair.

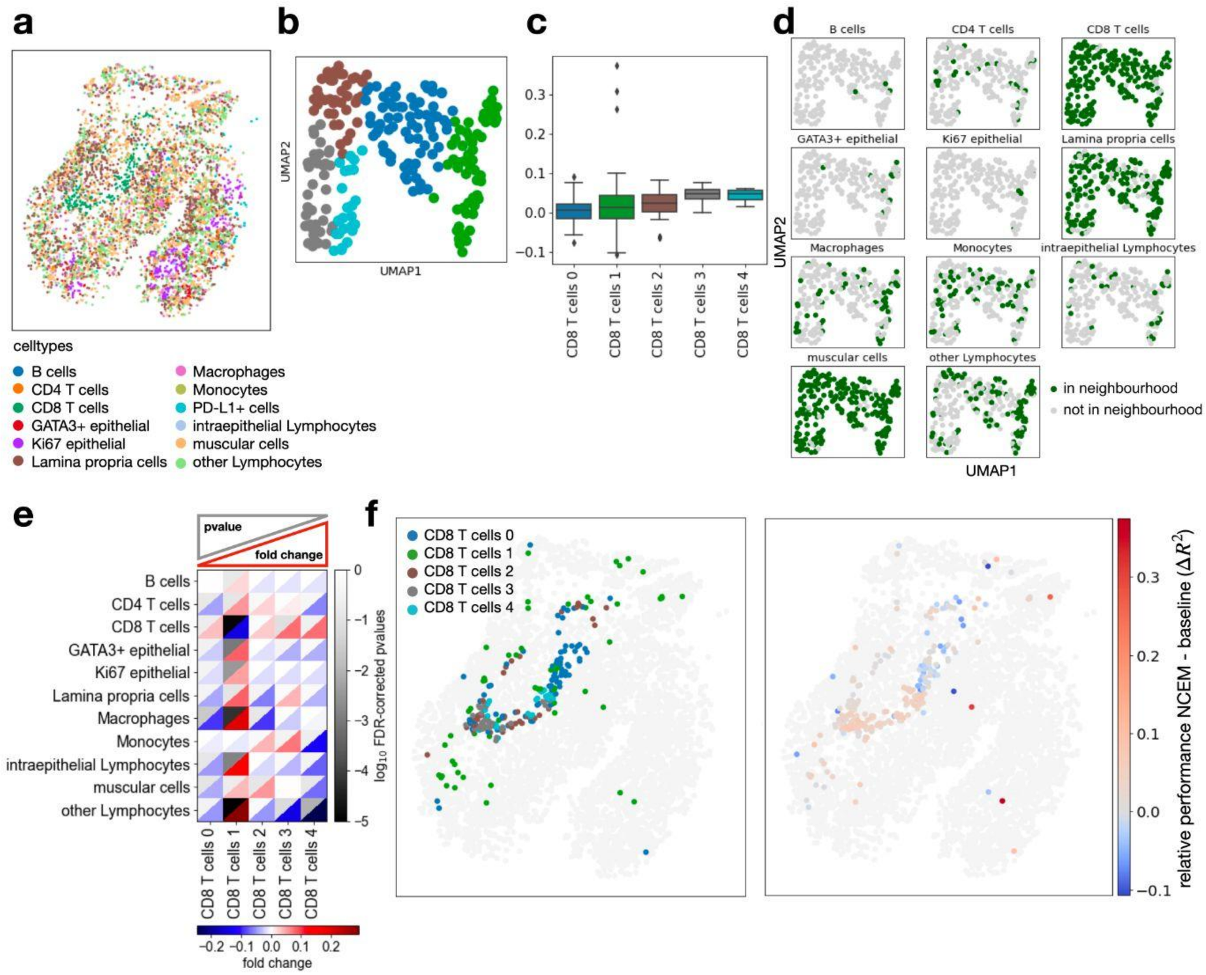

**Supp. Fig. 8: Attributing cell heterogeneity to niche composition in CD8 Tcells in inflamed colon. (a)** Area 1 of chip cytometry – colon dataset with the spatial allocation of all cell-types superimposed. **(b)** UMAPs of molecular embedding of CD8 T cells only with molecular sub-clustering superimposed (colors as in c)**. (c)** Distribution of cell-wise difference of $R^2$ between spatial model non non-spatial baseline model by molecular sub-cluster (CD8 T cells 0: n = 74, CD8 T cells 1: n = 58, CD8 T cells 2: n = 41, CD8 T cells 3: n = 37, CD8 T cells 4: n = 24). The centerline of the boxplots defines the mean over all relative $R^2$ values, the height of the box is given by the interquartile range (IQR), the whiskers are given by 1.5 * IQR and outliers are given as points beyond the minimum or maximum whisker. **(d)** UMAPs of molecular embedding of all CD8 T cells in area 1 (n = 234 cells) showing if a given cell-type is present in the neighborhood. The underlying neighborhoods were defined at the best performing resolution identified in Fig. 1c (40 μm). **(e)** Heatmap of fold change versus false-discovery rate corrected p-values of cluster enrichment of binary neighborhood labels, where fold changes are the ratio between the relative neighboring source cell-type frequencies per subtype cluster and the overall source cell-type frequency in the image. **(f)** Spatial allocation of area 1 of colon in the chip cytometry – colon dataset with CD8 T cell sub-states superimposed and superimposed the difference of $R^2$ between the NCEM interaction model at resolution of 40 μm and the best nonspatial baseline model.

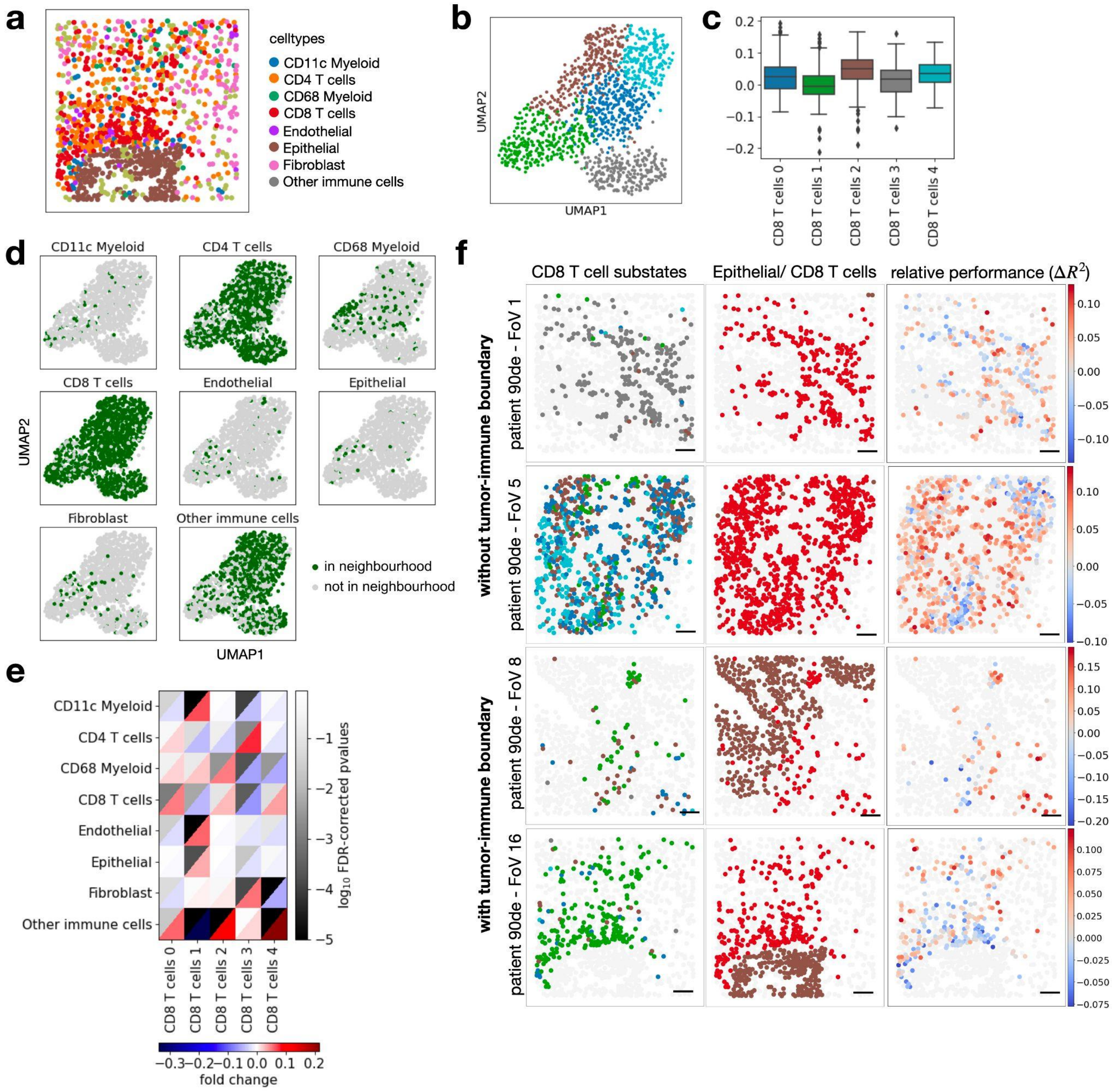


**Supp. Fig. 9: Attributing cell heterogeneity to niche composition in CD8 Tcells in colorectal cancer. (a)** Field of view 16 of MIBI TOF – cancer dataset with the spatial allocation of all cell-types superimposed. **(b)** UMAPs of molecular embedding of CD8 T cells only with molecular sub-clustering superimposed (colors as in c)**.** **(c)** Distribution of cell-wise difference of $R^2$ between spatial model non non-spatial baseline model by molecular sub-cluster (CD8 T cells 0: n = 304, CD8 T cells 1: n = 293, CD8 T cells 2: n = 278, CD8 T cells 3: n = 247, CD8 T cells 4: n = 207). The centerline of the boxplots defines the mean over all relative $R^2$ values, the height of the box is given by the interquartile range (IQR), the whiskers are given by 1.5 * IQR and outliers are given as points beyond the minimum or maximum whisker. **(d)** UMAPs of molecular embedding of all CD8 T cells in area 1 (n = 1,329 cells) showing if a given cell-type is present in the neighborhood. The underlying neighborhoods were defined at the optimal resolution identified in Fig. 1d (13 µm). **(e)** Heatmap of fold change versus false-discovery rate corrected p-values of cluster enrichment of binary neighborhood labels, where fold changes are the ratio between the relative neighboring source cell-type frequencies per subtype cluster and the overall source cell-type frequency in the image. **(f)** Spatial allocation of field of view 1, 5, 8 and 16 of colon in the MIBI TOF – cancer dataset with CD8 T cell sub-states superimposed and superimposed the difference of $R^2$ between the NCEM interaction model at a resolution of 13 µm and the best nonspatial baseline model (scale bar 50 µm).

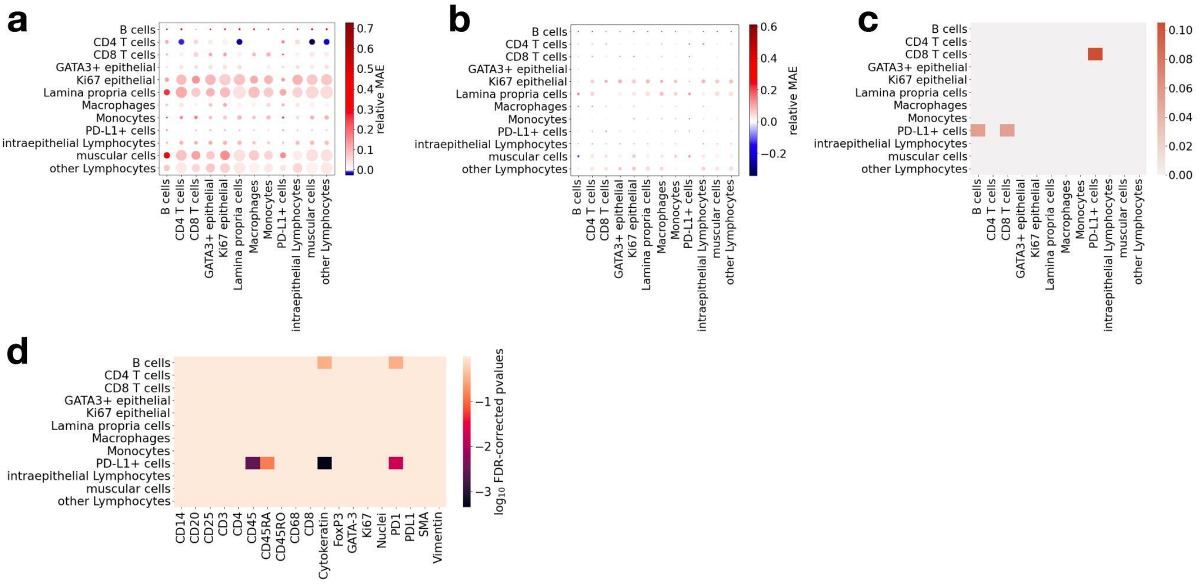


**Supp. Fig. 10: Parameter significance for chip cytometry – colon dataset. (a,b)** Relative predictive performance in terms of mean absolute error for the chip cytometry – colon dataset at 100 µm on training **(a)** and test data **(b)**. **(c)** Fraction of significant FDR-corrected p-values per sender–receiver cell-type pair for maximal euclidean edge length of 40 µm in chip cytometry – colon dataset. **(d)** log10 FDR-corrected p-values per gene for sender cell-type CD8 T cells for maximal euclidean edge length of 40 µm in chip cytometry – colon dataset.

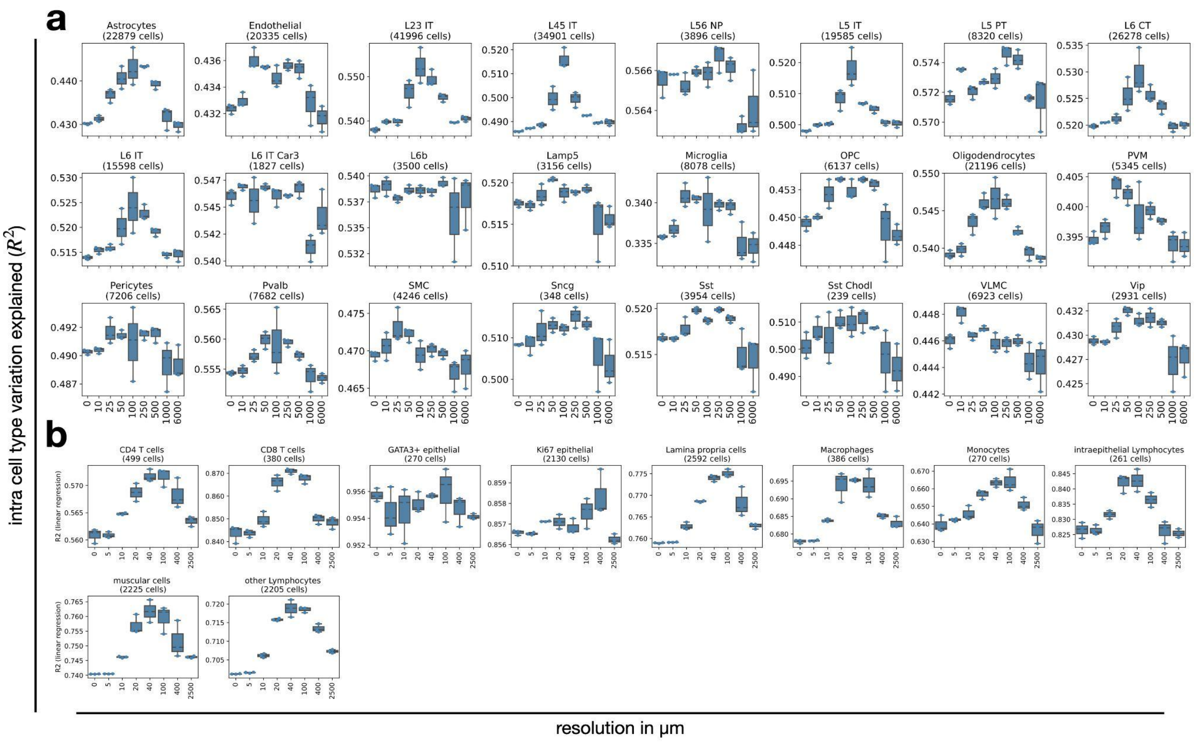


**Supp. Fig. 11: Length scales of dependencies for different target cell-types.** Shown are $R^2$ for held-out test data of ED-NCEM by resolution in µm for different predicted cell-types for MERFISH – brain data **(a)**, and chip cytometry – colon data **(b)** where each boxplot corresponds to a three-fold cross validation. For each box in **(a,b)**, the centerline defines the mean over all three cross validations, the height of the box is given by the interquartile range (IQR), the whiskers are given by 1.5 * IQR and outliers are given as points beyond the minimum or maximum whisker.

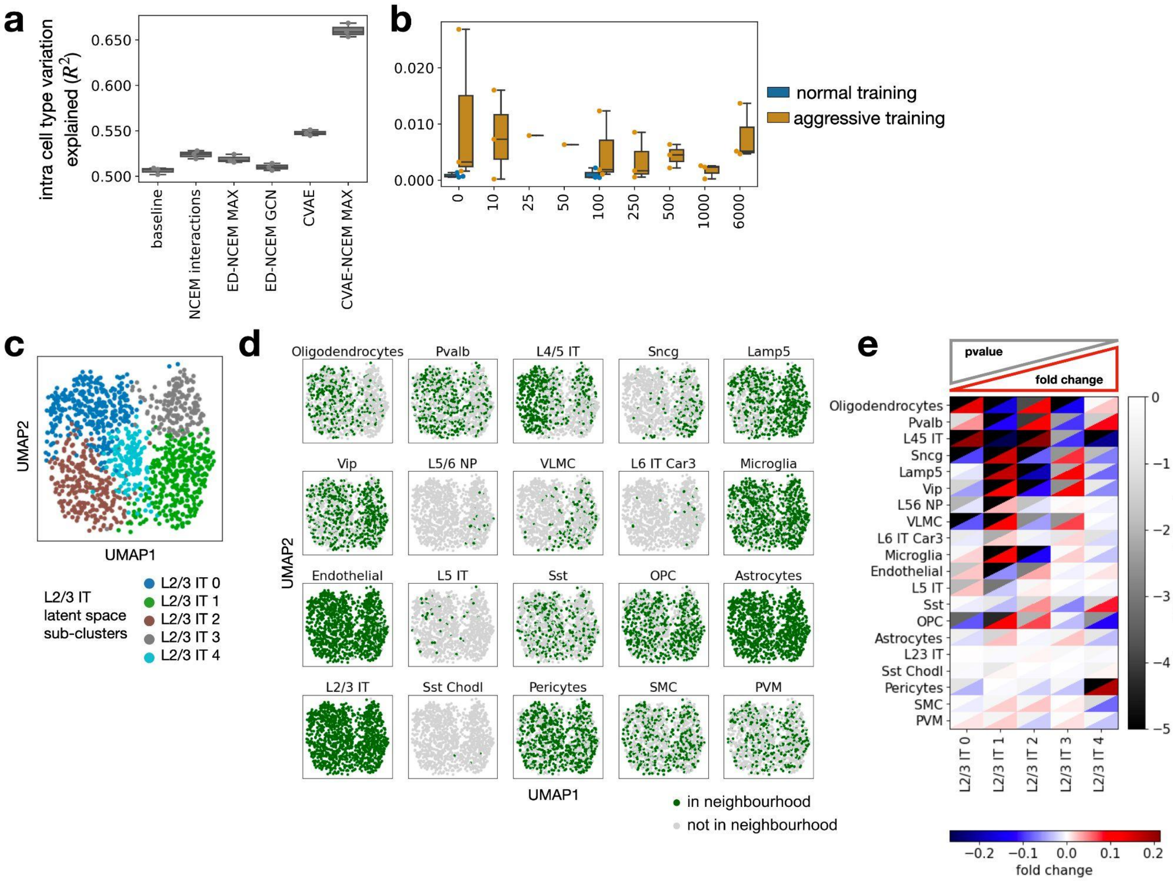


**Supp. Fig. 12: CVAE–NCEMs on MERFISH – brain data. (a)** Latent variable models improve reconstructive performance. Shown is the R2 of held-out test data based on forward pass model evaluation from MERFISH – brain data for linear models, encoder–decoder models and variational autoencoders for both non-spatial and spatial models. *baseline*: a nonspatial linear model of gene expression per cell-type; *NCEM interactions*: linear model with interaction effects; *ED*: encoder–decoder model; *IND*: the graph convolution is an indicator function across cell-types in the neighborhood; *GCN*: the graph convolution is a linear embedding (filter) of the cell-types in the neighborhood. **(b)** Neighborhood transfer performance of spatial and non-spatial models. Shown are the $R^2$ values of cells in the test set for models trained on predicting L2/3 IT cells for both CVAE models CVAE–NCEMs trained on neighborhoods with different radii with optimization algorithm as color. *Plain*: normal CVAE training; *aggressive*: aggressive encoder training. **(c-e)** Latent variables of CVAE–NCEM are confounded with neighborhood conditions. **(c)** UMAP of molecular embedding in the CVAE–NCEM IND latent space of L2/3 IT cells in an example image (n = 1204 cells) with molecular sub-clustering superimposed (L2/3 IT 0: n = 323, L2/3 IT 1: n = 315, L2/3 IT 2: n = 250, L2/3 IT 3: n = 170, L2/3 IT 4: n = 146)**.** **(e)** UMAPs of molecular embedding in the CVAE–NCEM IND latent space of all L2/3 IT cells in the same image with superimposed binary label of presence of a given cell-type, as defined in the title, in the neighborhood. The underlying neighborhoods were defined at a resolution of 100 µm. **(f)** Heatmap of fold change versus false-discovery rate corrected p-values of cluster enrichment of binary neighborhood labels where fold changes are the ratio between the relative neighboring source cell-type frequencies per subtype cluster and the overall source cell-type frequency in the image.

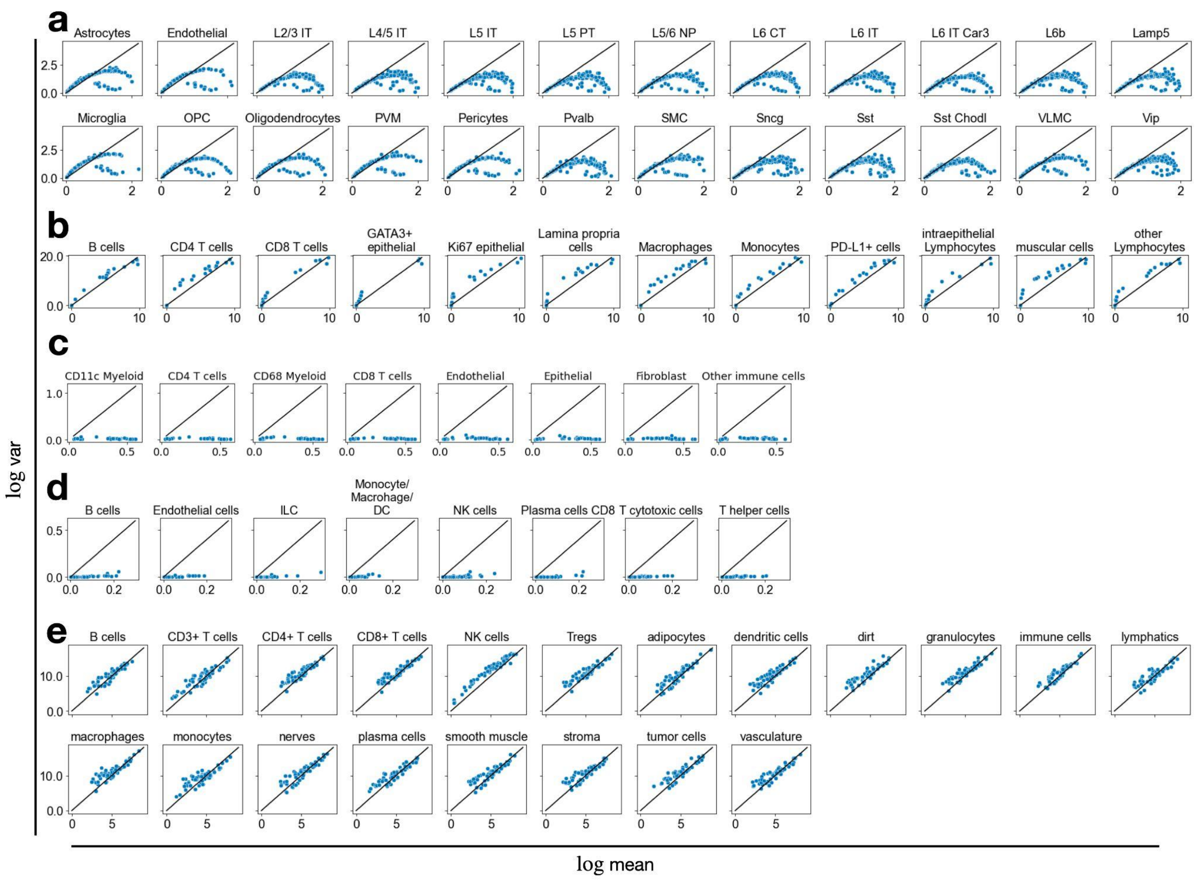


**Supp. Fig. 13: Distributional characteristics of gene expression measurements of single cells from spatial molecular profiling assays.** Shown is the mean variance plot over observed genes for MERFISH – brain data **(a)**, chip cytometry – colon data **(b)**, MIBI TOF – cancer data **(c)**, MELC – tonsils data **(d)** and CODEX – cancer data **(e)**.
